# Dissecting the dynamics of signaling events in the BMP, WNT, and NODAL cascade during self-organized fate patterning in human gastruloids

**DOI:** 10.1101/440164

**Authors:** Sapna Chhabra, Lizhong Liu, Ryan Goh, Aryeh Warmflash

**Affiliations:** Systems, Synthetic, and Physical Biology graduate program, Rice University, Houston TX 77005; Department of Biosciences, Rice University, Houston TX 77005; Department of Mathematics, Boston University, Boston, MA 02215; Department of Bioengineering, Rice University, Houston TX 77005

**Author notes:** Correspondence to AW.

**Keywords:** Gastrulation, human embryonic stem cells, signaling dynamics, fate patterning

## Abstract

During gastrulation, the pluripotent epiblast is patterned into the three germ layers, which form the embryo proper. This patterning requires a signaling cascade involving the BMP, WNT and NODAL pathways; however, how these pathways regulate one another in space and time to generate cell-fate patterns remains unknown. Using a human gastruloid model, we show that BMP signaling initiates a wave of WNT signaling, which, in turn, initiates a wave of NODAL signaling. While WNT propagation depends on continuous BMP activity, NODAL propagates independently of upstream signals. We further show that the duration of BMP signaling determines the position of mesodermal differentiation while WNT and NODAL synergize to achieve maximal differentiation. The waves of both WNT and NODAL signaling activity extend farther into the colony than mesodermal differentiation. Combining dynamic measurements of signaling activity with mathematical modeling revealed that the formation of signaling waves is inconsistent with WNT and NODAL forming a stable spatial pattern in signaling activities, and the final signaling state is spatially homogeneous. Thus, dynamic events in the BMP, WNT, and NODAL signaling cascade, in the absence of a signaling gradient, have the potential to mediate epiblast patterning.

## Introduction

Gastrulation is a crucial stage in embryonic development when a homogeneous population of pluripotent epiblast cells self-organizes to form the three germ layers: endoderm, mesoderm and ectoderm, which develop into the embryo. Insights into mammalian gastrulation come from decades of genetic and biochemical studies in the mouse embryo (1). These studies have revealed that a signaling cascade involving the BMP, WNT and NODAL pathways is integral for initiating gastrulation. BMP signaling in the extra-embryonic ectoderm activates WNT signaling in the epiblast and the overlying visceral endoderm (2,3). WNT signaling activates NODAL signaling in these two tissues and NODAL signaling, in turn, feeds back to maintain BMP signaling in the extra-embryonic ectoderm (3,4). This signaling feedback between BMP, WNT and NODAL pathways initiates the formation of primitive streak that marks the onset of gastrulation. Although the signaling pathways involved are known, their activities in space and time, regulation and coordination with other cellular processes, like cell division and migration, during epiblast patterning are largely unknown (1).

In a previous study, we showed that spatially confined human embryonic stem cells (hESCs), treated with BMP4 (gastruloids) self-organize to form radial patterns of distinct germ layers: an outer ring of extra-embryonic cells, followed by endodermal and mesodermal rings, and an ectodermal center, thus recapitulating some aspects of gastrulation *in vitro* (5). These findings have since been reproduced in other labs (6), and a comparable system for mouse ESCs has been developed (7). Three-dimensional models have also been developed that recapitulate aspects of early mammalian development (8–11), and in some cases, even morphologically resemble embryos (12–14). Although the germ layer patterns in gastruloids differ from the trilaminar germ layer patterns *in vivo*, they offer a reproducible, quantitative, and controlled system to understand mechanisms underlying germ layer patterning.

Recent studies have provided insights into the mechanisms underlying the self-organized fate patterning of gastruloids. Exogenous BMP4 initially activates signaling homogeneously throughout the colony, but within 12h, signaling is restricted to the colony edges (15). This restriction is caused by upregulation of BMP inhibitor Noggin together with a lack of BMP receptor accessibility at the center (16). Prolonged BMP signaling at the colony edge leads to differentiation towards an extra-embryonic fate (17). In response to BMP treatment, hESCs activate paracrine WNT and NODAL signaling, which are both necessary for mesendodermal ring formation (5,18). In contrast to BMP signaling, NODAL signaling activity is highly dynamic: a wave of signaling moves from the edge towards the colony center at a constant rate, specifying the mesendodermal rings in its wake ((15), Figure S1E). However, the dynamics of interactions between the BMP, WNT and NODAL signaling pathways and the features of signaling that govern cell-fate patterning are not understood.

In this study, we quantitatively examined the temporal requirement of signals at each step in the BMP-> WNT-> NODAL signaling cascade for the dynamics of downstream signals and the subsequent self-organized fate patterning of gastruloids. Our results reveal that BMP signaling initiates and maintains a wave of paracrine WNT signaling, which moves towards the colony center at a constant rate. WNT initiates a wave of paracrine NODAL signaling, which, unlike WNT, moves inwards independently of upstream signaling. We further show that the duration of BMP signaling determines the position of mesoderm differentiation while WNT and NODAL synergize to achieve maximal mesoderm differentiation. The waves of both WNT and NODAL signaling activities extend further towards the center of colony than the region of mesodermal differentiation, suggesting that cell-fate patterning is not a result of a spatial signaling pattern. A simple reaction-diffusion based mathematical model comprised of an activator-inhibitor system revealed that the final signaling state can be explained by the tendency towards a homogenous signaling state combined with boundary effects. There is no stable spatial pattern of WNT and NODAL signaling activities that underlies fate patterning. Taken together, our data suggest that the dynamics of signaling events in the cascade involving BMP, WNT and NODAL pathways, and not a spatial pattern in signaling, controls the self-organized fate patterning of human gastruloids.

## Results

### WNT signaling initiates and NODAL signaling upregulates mesodermal differentiation

To examine the role of paracrine signals secreted by cells without influence from undefined components present in mouse embryonic fibroblast conditioned media (MEF-CM), we performed the micropatterned gastrulation assay in the defined mTeSR1 media as outlined in (19). In mTeSR1, hESCs treated with BMP4 self-organize to form an outer ring of CDX2+ extra-embryonic cells, and an inner ring of BRACHYURY (BRA+) primitive streak or mesodermal cells (Figure 1A). SOX17+ endodermal cells fall in between these two (FigureS1A), as observed previously in MEF-CM (5). Inside of the BRA+ ring, center cells co-express the pluripotency markers NANOG and SOX2, with the centermost cells showing lower NANOG levels (Figure S1B), a possible sign of the onset of ectodermal differentiation (20). Compared to the MEF-CM protocol in which NANOG expression was completely lost in the center cells (5), the mTeSR1 protocol recapitulates an earlier time point in gastrulation when primitive streak formation has begun, but the remainder of the epiblast remains pluripotent with only shallow gradients in pluripotency markers such as NANOG (1,7).

**Figure 1:**
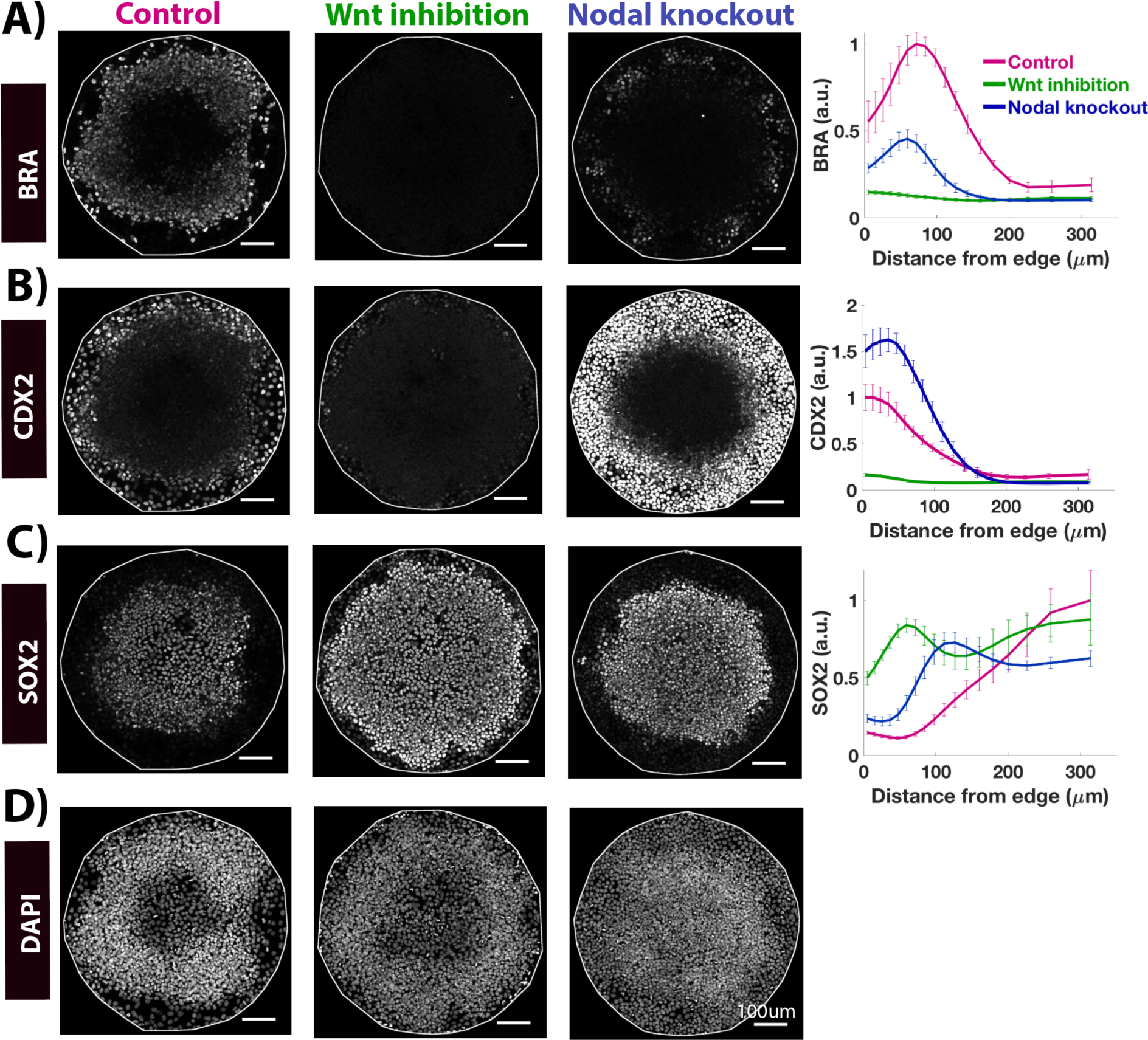
Paracrine WNT and NODAL signaling are necessary for the self-organized differentiation of micropatterned hESCs. **(A-D)** Images of samples immunostained for the indicated markers in different conditions: Control and NODAL^-/-^ cells were treated with 50ng/ml BMP4. WNT inhibition refers to wildtype cells treated with 50ng/ml BMP and 5 µM IWP2. All samples were fixed at 44 post BMP4 treatment. Quantification represents intensity levels of indicated markers normalized to DAPI, averaged at different positions along the colony radii. Error bars represent standard error of the mean. N >= 10. Scale bar =100µm. Colony diameter = 700µm. See also Figures S1, S2, S7.

To determine the role of paracrine WNT and NODAL signaling in this self-organized differentiation, we first inhibited these signals by chemical and genetic perturbations. To inhibit paracrine WNT signaling, we added IWP2, which inhibits the secretion of all WNT ligands (21), at the beginning of the gastrulation assay. To inhibit paracrine NODAL signaling, we created NODAL knockout cells (NODAL-/-) using CRISPR-Cas9 and used these cells in the gastrulation assay. Although these cells remain pluripotent by responding to exogenous NODAL pathway signals in media, they do not produce functional NODAL protein (Figure S2), thus specifically inhibiting paracrine NODAL signaling.

Studies in various model organisms and *in vitro* systems have established the necessity of WNT and NODAL signaling for the formation of primitive streak, where mesoderm and endoderm cells originate (1,22). Consistent with these, removing either paracrine WNT or NODAL signaling reduced the number of BRA+ mesodermal cells. This loss was complete in the case of WNT inhibition (Figure 1A). In addition to NODAL secreted by cells, TGFβ1 present in mTeSR1 also activates the NODAL signaling pathway. Inhibiting the NODAL pathway activity downstream of ligand-receptor binding using a small molecule inhibitor of receptor kinase activity, SB431542 (23) (SB), had a more severe effect on mesodermal differentiation, as shown previously ((5,18), Figure S1C,D). Consistent with this, NODAL knockout cells show high nuclear SMAD2 - a signal transducer of active NODAL/TGF-β signaling, at the colony edges, which is lost upon treatment with SB (Figure S1C). However, even complete inhibition of NODAL signaling does not result in complete loss of BRA+ cells as observed upon inhibition of WNT signaling. This suggests that paracrine WNT signaling is necessary and sufficient to initiate mesoderm differentiation, whereas paracrine NODAL signaling increases the fraction of cells differentiating towards mesoderm. Endoderm differentiation, on the other hand, requires both WNT and NODAL signaling for its initiation as inhibition of either pathway completely abolished SOX17 expression (Figure S1A,D).

### WNT signaling upregulates and NODAL signaling downregulates extra-embryonic differentiation

Surprisingly, WNT inhibition almost completely abolished CDX2+ cells at the colony edges (Figure 1B), suggesting that paracrine WNT signaling is required for this fate in addition to the BRA+ streak fates. This suggests that these CDX2+ cells might represent late-streak extra-embryonic mesodermal cells, which originate in the posterior epiblast of the mouse embryo where WNT signaling is high (1). Alternatively, it points to a previously unappreciated role of WNT signaling for trophectoderm differentiation (24,25). Interestingly, although WNT inhibition prevents differentiation of both extraembryonic and mesendoermal cells, the expansion of pluripotent SOX2+ cells does not reach the colony edges (Figure 1C). Instead, there exists a population of cells at the colony edges that that do not express these differentiation markers or those of pluripotency. These results raise the questions of the identity of the CDX2+ cells at the colony border and the cells that result in this location from the inhibition of WNT signaling.

Although WNT signaling is necessary for CDX2+ differentiation, WNT treatment alone is insufficient to induce CDX2 expression in micropatterned hESCs (18). Thus, WNT, in combination with BMP4 signaling instructs CDX2+ expression in cells at the colony edges. In the absence of paracrine NODAL signaling, cells preferentially respond to exogenous BMP4 and paracrine WNT signaling by upregulating CDX2 as indicated by higher average levels and a broader domain (Figure 1B). Taken together, these results show that paracrine WNT and NODAL signaling are both necessary for self-organized differentiation of micropatterned hESCs, but the effects of removing them are different, especially with regards to extra-embryonic differentiation at the colony periphery.

### The domain of active WNT signaling expands inwards during patterning

To better understand the role of paracrine WNT signaling in spatial patterning, we used our previously developed transgenic hESC line in which GFP is inserted into the endogenous β-catenin locus to form a fusion protein (hereafter referred to as GFP-β-cat hESCs, (26)). β-catenin is stabilized by WNT signaling and serves as a transcription factor to activate WNT pathway targets. However, β-catenin also strengthens cell-cell junctions at the cell membrane. Thus, to avoid misinterpreting signaling activity due to the membrane population of β-catenin, we considered only non-membrane fluorescence as a measure of WNT signaling activity (Figure S7).

We examined the dynamics of WNT signaling during self-organized differentiation using time-lapse imaging of GFP-β-cat hESCs seeded onto micropatterned colonies, from 3 to 47h post BMP4 treatment (Figure 2A, Movie S1). In the first 20h following treatment, the colonies initially contract in response to withdrawal of Rock-Inhibitor and then spread to fill the entire micropatterned space available. During this time, WNT signaling increases throughout the colony, with the colony edges showing slightly higher signaling than the rest. From 20-40h, the peak signaling increases continuously, but from 40-47h, it decreases to a slightly lower value (Figure 2B, S3A).

**Figure 2:**
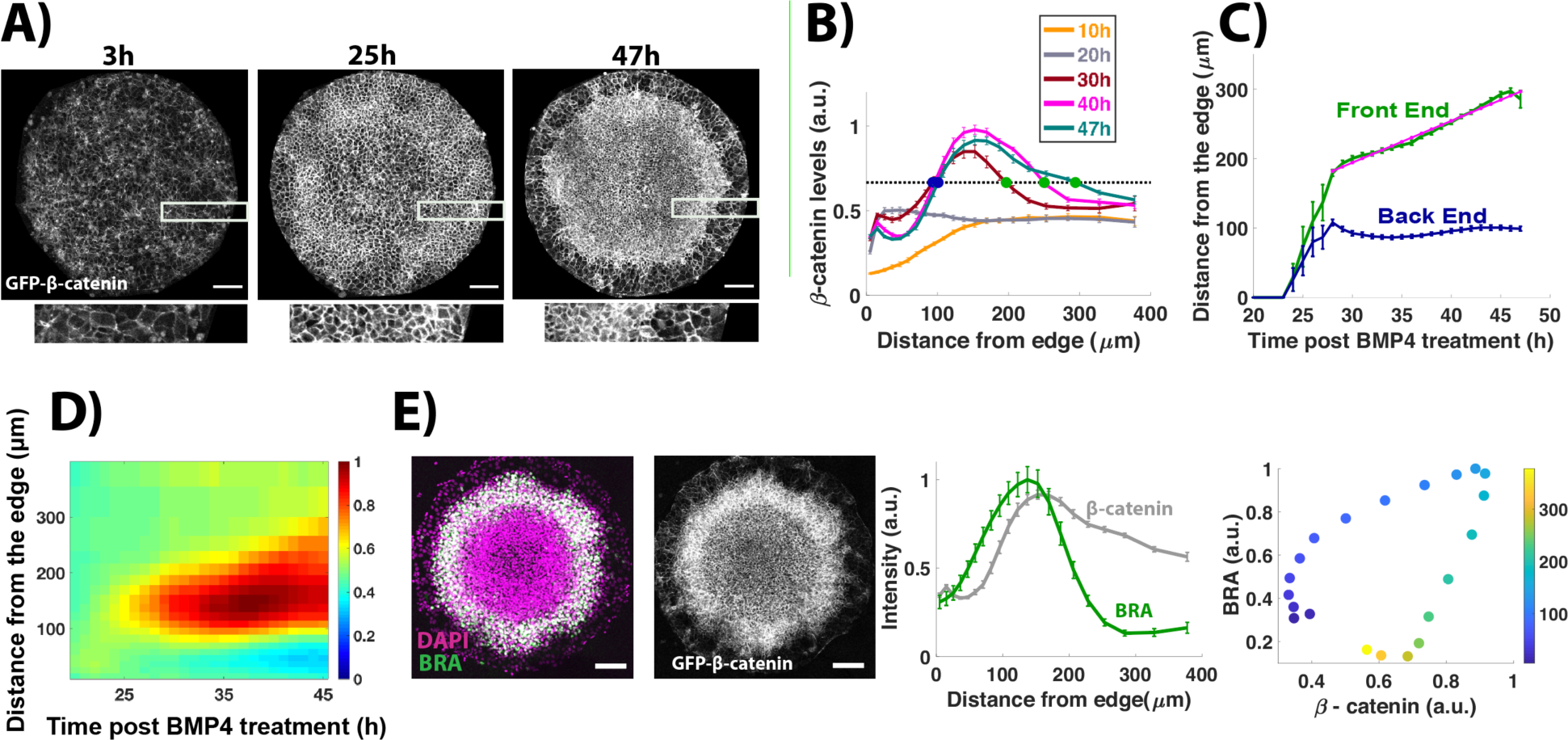
The domain of active WNT signaling expands inwards during patterning. **(A)** Snapshots of GFP-β-catenin hESCs from time-lapse imaging at indicated time points post BMP treatment. Marked regions are magnified in images shown below. **(B)** Average non-membrane β-catenin intensity levels as a function of distance from the colony edge. The legend indicates the time post BMP treatment represented by each curve. Green and blue dots represent the front and back edges of the signaling trajectory, respectively. **(C)** The positions of the front and back of the signaling domain as a function of time post BMP addition. Magenta line shows linear fit with equation 6.02 µm/h *t + 13.57 µm, R^2^= 0.98. **(D)** Kymograph showing spatiotemporal evolution of non membrane β-catenin levels. At time points earlier than the first time plotted in D, signaling is below threshold signaling at all positions in the colony. **(E)** Image of a colony immunostained for BRA and DAPI following time-lapse imaging. Snapshot from time lapse imaging for the same colony at 47h post BMP treatment. (Left) Quantification represents average intensity levels for BRA and non-membrane β-catenin, normalized by DAPI, plotted as a function of radial position. Error bars represent standard error of the mean. N= 9. (Right) Average BRA intensity levels as a function of β-catenin levels, color-coded distance from the colony edge (µm). Scale bar: 100µm. See also S3, S7, movieS1

We quantified the spatial dynamics of WNT signaling by defining a territory of active signaling as the region with non-membrane β-catenin levels greater than half the maximal WNT signaling in the entire time course, and traced the position of this territory in time (Figure S3B). We note that this definition is a convenient way to quantify the spatial extent of WNT signaling, but none of our conclusions depend on the threshold for active signaling. There is no active WNT signaling in the first 24h post BMP4 treatment. From 24-27h, active signaling occurs near the colony edges in some colonies. From 27h onwards, the active signaling forms a ring-like domain whose inner edge (front end) moves towards the colony center at a constant rate, while the outer edge(back end) remains stationary, resulting in continuous expansion of the active WNT signaling domain (Figure 2C,D).

To determine the relationship between WNT signaling and mesodermal differentiation, we immunostained the colonies for BRA following live-cell imaging. Although the territories of active WNT signaling and BRA overlap, WNT signaling extends further towards the colony center than the mesodermal ring (Figure 2E, S3D,E). There is no simple correspondence between mesodermal differentiation and WNT signaling levels. At all times during differentiation, the same level of WNT signaling on the inner and outer sides of the peak yield dramatically different levels of BRA (Figure S2E).

### Duration of BMP signaling controls the position of BRA+ mesodermal ring by modulating the position of peak WNT signaling

BMP is necessary to trigger the dynamic waves of WNT and NODAL that cause mesoderm differentiation, but whether ongoing BMP signaling is necessary for the propagation of these waves remains unclear. To test this, we added a BMP4 signaling inhibitor, LDN193189 (27) (LDN), at different time points during the gastrulation assay and quantified the resultant WNT and NODAL signaling and cell fates. We verified the function of LDN by immunostaining for activated SMAD1/5/8, the key signal transducers for BMP signaling (Figure S4A).

To determine the effect of BMP inhibition on NODAL signaling, we immunostained the LDN treated samples for SMAD2, a signal transducer that enters the nucleus in response to NODAL signaling. When BMP4 signaling is allowed for the entire 48h of the gastrulation assay, a wave of SMAD2 moves from colony edge inwards, eventually reaching the colony center (Figure S1C). Upon inhibition of BMP signaling 10h post treatment, the front of active SMAD2 stops halfway to the center, but BMP signaling inhibition at or beyond 15h does not prevent active SMAD2 from reaching the colony center, indicating that continuous BMP4 signaling is not required for the inward movement of NODAL signaling. Interestingly, when LDN was added at 15h it strengthened the NODAL signaling at the colony border, indicating that the continuous BMP signaling at the border downregulates NODAL signaling and this likely plays a role in differentiation to CDX2+ extraembryonic fates (see below). Thus, 15h of BMP signaling are sufficient for maximal activation of NODAL, and further BMP signaling primarily downregulates NODAL at the colony edge.

WNT signaling serves as an intermediary between BMP and NODAL (28). It is possible that WNT activity movement is similarly independent of continued BMP signaling, or alternatively, NODAL may move inward independently of WNT. To test these hypotheses, we added LDN during live cell imaging of micropatterned GFP-β-cat hESCs at three different time points (0h, 11h, 23h) which correspond to the negative control which removes all BMP signaling (0h) and the time points when the NODAL wave has either moved halfway (11h) or the entire way to the colony center (23h). Quantification of WNT signaling dynamics shows that even in the sample treated with LDN at the latest time point (23h), after the movement of NODAL signaling becomes independent of BMP signaling, WNT signaling settles to a lower peak value and the domain of active signaling does not travel as far. (Figure 3A-D, S4A, movie S2-S5). The peak WNT signaling activity is also restricted to a territory closer to the colony edges, suggesting that continuous BMP signaling is necessary for the inward shift of WNT signaling activity. This is in contrast to NODAL signaling that travels to the center of the colony when BMP signaling is terminated after 15h, suggesting that following its initiation, the NODAL signaling wave propagates independently of WNT signaling.

**Figure 3:**
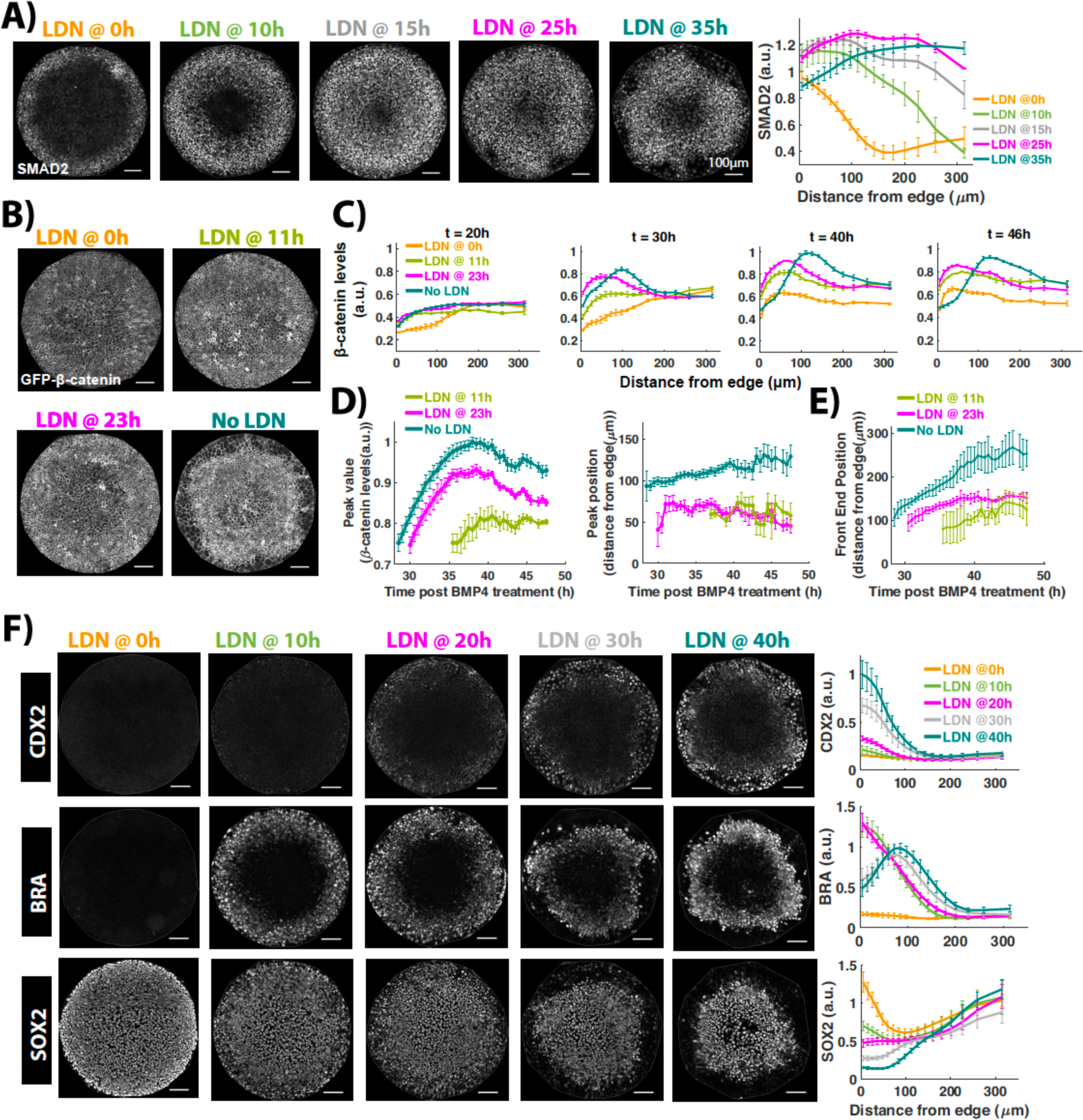
Duration of BMP signaling controls the position of BRA+ mesodermal ring by modulating the position of peak WNT signaling. **(A)** (left) Images of samples immunostained for SMAD2 after 44h of BMP treatment. The time between BMP4 and 200nM LDN addition is indicated above each image. (right) Quantification represents average SMAD2 nuclear intensities normalized to DAPI as a function of radial position. Error bars represent standard error. N>=5. **(B)** Snapshots of GFP-β-catenin hESCs from time-lapse imaging movies at 46h post BMP treatment. Time between BMP4 and LDN addition is indicated above image. Error bars represent standard error. N=3 for LDN@0h, LDN@11h, for others N>=5. **(C)** Average non-membrane β-catenin levels as a function of radial position. The timing of LDN addition after BMP4 treatment for each curve is shown in the legend while the time being analyzed in shown above the plot. **(D)** Temporal evolution of the position and intensity of peak signaling (defined by maximal non-membrane β-Catenin intensity). **(E)** Temporal evolution of the front of the domain of active signaling. In (D,E) at time points earlier than the first one in each curve, signaling was below threshold signaling at all positions. **(F)** Images of samples immunostained for CDX2, BRA, SOX2 after 44h of BMP treatment. The time between BMP4 and LDN addition is indicated above the image. Quantification represents average nuclear intensities of indicated markers normalized to DAPI as a function of radial position. Error bars represent standard error. N>=10. Scale bar = 100um. See also S4, S7, movie S2,S3,S4,S5.

To determine the functional consequence of modulating the duration of BMP signaling on cell fates, we immunostained the LDN treated samples for the fate markers CDX2, BRA, and SOX2. Inhibition of BMP signaling in the first 20h leads to an outward shift of fate patterns, with a loss of CDX2+ cells and the presence BRA+ cells at the colony edges. Allowing BMP signaling for 30h or more yields CDX2+ cells at the colony edges, followed by a BRA+ inner ring, suggesting that continuous BMP signaling for at least 30h is necessary for CDX2+ differentiation at colony edges (Figure 3F) in the absence of which edge cells differentiate towards mesodermal fate.

### Duration of WNT secretion determines peak WNT signaling levels

To determine whether continuous WNT secretion is required for inward propagation of the WNT signaling, we inhibited WNT secretion using IWP2 during differentiation of micropatterned GFP-β-cat hESCs, before (15h) and after (30h) WNT signaling first peaks in the future mesodermal ring (25h, Figure S3A), and recorded the resulting WNT signaling by time-lapse imaging.

Inhibition of WNT secretion lowers the peak WNT signaling levels in a time-dependent manner (Figure 4A-D), with inhibition at the earliest time point giving the lowest peak signaling. Surprisingly, despite a reduction in signaling levels, the spatial dynamics of WNT signaling in the colony are similar: WNT activity first increases in a ring-like domain and then moves inwards towards the colony center (Figure 4D, E, S4C-D, movieS6-S8). The inward movement happens at the same rate in all conditions as indicated by the movement of WNT signaling fronts (Figure 4E). This suggests that the movement of the wave of WNT signaling is independent of secretion of new WNT proteins and may rely on diffusive or active extracellular movement of the WNT proteins produced near the colony edge. In contrast, WNT secretion is required to increase the total levels, but not the spatial extent, of WNT activity.

**Figure 4:**
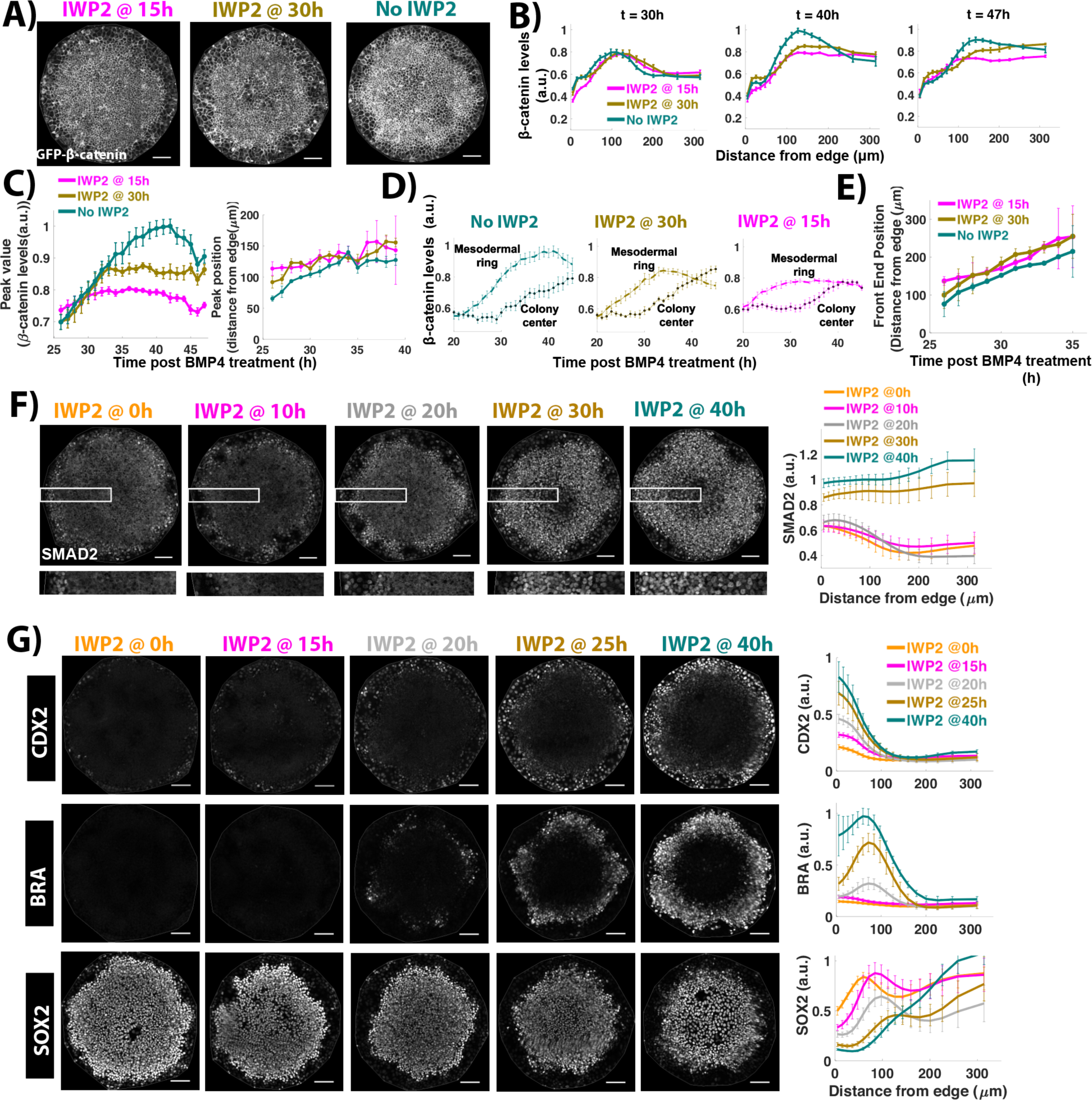
Continuous WNT secretion is necessary to achieve high WNT signaling and BRA+, CDX2+ differentiation levels. **(A)** Snapshots of GFP-β-catenin hESCs from time-lapse imaging at 46h post BMP treatment. Time between BMP4 and IWP2 addition indicated above each image. Error bars represent standard error. N = 5. **(B)** Average non-membrane β-catenin levels as a function of radial position. The timing of IWP2 addition after BMP treatment is shown in the legend while the time being analyzed is shown above the plot. **(C)** Temporal evolution of the position and intensity of peak signaling (defined by maximum non-membrane β-catenin intensity). **(D)** Temporal evolution of average non-membrane β-catenin levels in a ring inside the region of mesodermal differentiation (distance from edge: 134.13-150.75µm) and at the center (distance from edge: 277.89-350µm) of the colony. **(E)** Temporal evolution of the front of the domain of active signaling. (In C&E, at time points earlier than the first one in each curve, signaling was below threshold signaling. In C, the last time point indicates time when signaling curve flattens for sample treated with IWP2@15h. In E, the last time point indicates time when the entire signaling front is higher than the threshold). **(F)** Images of samples immunostained for SMAD2 after 44h BMP treatment. Time of IWP2 addition after BMP treatment is indicated above image. Quantification represents average nuclear intensities normalized to DAPI as a function of radial position. Bottom row: high magnification view of the rectangular region marked in images above. N>=5. **(G)** Images of samples immunostained for CDX2, BRA, SOX2 after 44h of BMP treatment. The time between BMP4 and IWP2 addition is indicated above the image. Quantification represents average nuclear intensities of indicated markers normalized to DAPI as a function of radial position. Error bars represent standard error. N>=10. Scalebar = 100um. See also Figure S4, movie 6,7,8.

### Duration of WNT secretion controls inward movement of NODAL signaling activity and the levels of BRA+ mesoderm and CDX2+ extra-embryonic differentiation

To determine the functional consequence of reduced WNT signaling levels, we inhibited WNT secretion at different time points and examined the effect on initiation of paracrine NODAL signaling and the differentiation of BRA+ mesodermal and CDX2+ extra-embryonic fates all of which depend on WNT signaling.

The data from the BMP inhibition experiments above show that at some point, the movement of the WNT and NODAL waves become independent (Figure 3A-E). To determine when this occurs, we immunostained the IWP2 treated samples for SMAD2/3. We observed a binary effect on the movement of NODAL signaling. Inhibition in the first 20h completely prevents the inward movement of NODAL signaling and activity is restricted to the colony edges. NODAL signaling in these conditions is comparable to that observed in NODAL-/- cells indicating that early WNT inhibition abolishes the effects of paracrine NODAL (Figure 4F, S1C). Secretion inhibition at 30h and beyond has no effect on inward movement of NODAL signaling activity, while at 25h we observe a mixture of these two phenotypes (Figure S4B). Thus, the NODAL wave is initiated between 20 and 30h, consistent with our previous live cell measurements of signaling dynamics (15) and rapidly becomes independent of WNT signaling.

WNT secretion is necessary to initiate BRA expression in the inner ring, however, is it also necessary to achieve maximal mesodermal differentiation? Quantifying BRA levels after IWP2 addition at different times shows that 15-20h of WNT secretion is necessary to form BRA+ cells. Longer than 20h of WNT secretion increases the number of BRA+ cells widening the BRA+ domain, with the longest duration giving the widest BRA+ ring (Figure 4G), suggesting that continuous WNT secretion is necessary to increase the fraction of BRA+ cells in the mesodermal ring.

WNT and BMP4 signaling instruct CDX2+ differentiation in cells at colony edges and WNT secretion is necessary for CDX2+ differentiation (Figure 1), but is continuous secretion necessary to sustain CDX2+ differentiation? Varying the timing of IWP2 addition shows that longer durations of WNT secretion increase CDX2 levels at colony edges, with the longest duration giving the highest levels (Figure 4G), suggesting that continuous WNT secretion is necessary for high CDX2 expression in the edge cells. Given that WNT signaling activity consistently remains low in edge cells (Figure2), this result raises the question of how CDX2 expression is regulated by WNT signaling which requires further investigation.

### Duration of NODAL signaling determines the level of CDX2+ extra-embryonic differentiation and BRA+ mesodermal differentiation

Inhibition of NODAL signaling increases CDX2 levels in extraembryonic edge cells and lowers BRA levels in mesodermal cells, suggesting that during spatial patterning, NODAL signaling activity acts as a positive regulator of BRA+ mesodermal differentiation and a negative regulator of CDX2+ extra-embryonic differentiation. (Figure1, S1). Is continuous NODAL signaling required to achieve these effects? To examine this, we inhibited NODAL signaling at different time points during spatial patterning using SB and immunostained for markers of cell fates.

Inhibiting NODAL signaling within the first 15h had the same effect as the absence of NODAL signaling, with both conditions resulting in high CDX2+ levels in the outer ring and low BRA+ levels in the inner ring, compared to the control sample (Figure 5), suggesting that NODAL signaling in the first 15h of patterning has no effect on the final pattern. Increasing the duration of NODAL signaling decreases CDX2+ differentiation and increases BRA+ differentiation in a time-dependent manner, with the longest signaling duration having the lowest CDX2 levels and highest BRA+ levels (Figure 5), suggesting that continuous NODAL signaling is necessary to limit CDX2+ differentiation to the outer ring and achieve the highest levels of BRA+ differentiation in the inner ring.

**Figure 5:**
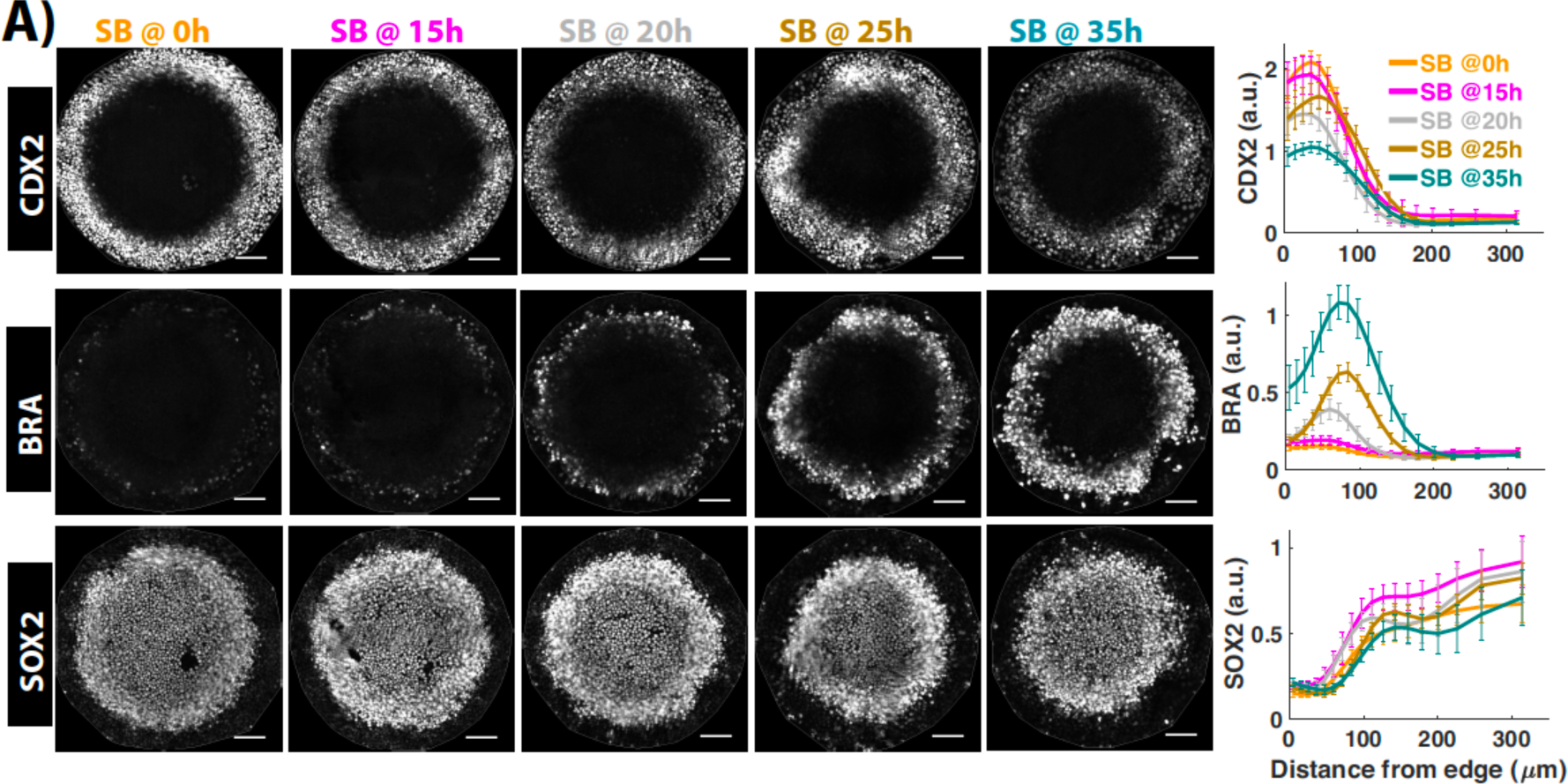
Continuous NODAL signaling is necessary to limit CDX2+ differentiation in outer ring and achieve high BRA+ differentiation in inner ring. **(A)** Images of samples immunostained for fate markers after 44h of BMP treatment. The time between BMP4 and SB addition is indicated above the image. Quantification represents average nuclear intensities of indicated markers normalized to DAPI as a function of radial position. Error bars represent standard error. N>=10. Scale bar = 100um

### Inward movement of signaling activities is not due to movement of cells

The inward movement of paracrine signaling activities during spatial patterning could be a result of cells with high signaling activity actively migrating inwards. To test this hypothesis, we tracked the movement of individual cells during self-organized differentiation. To improve tracking efficiency, we mixed cells labeled with an YFP-H2B fusion protein with unlabeled cells in the ratio 1:100 prior to seeding. Six colonies were imaged every 10 minutes during the time when the WNT signaling activity spreads inwards (Figure 6A,C, Movie S9).

**Figure 6:**
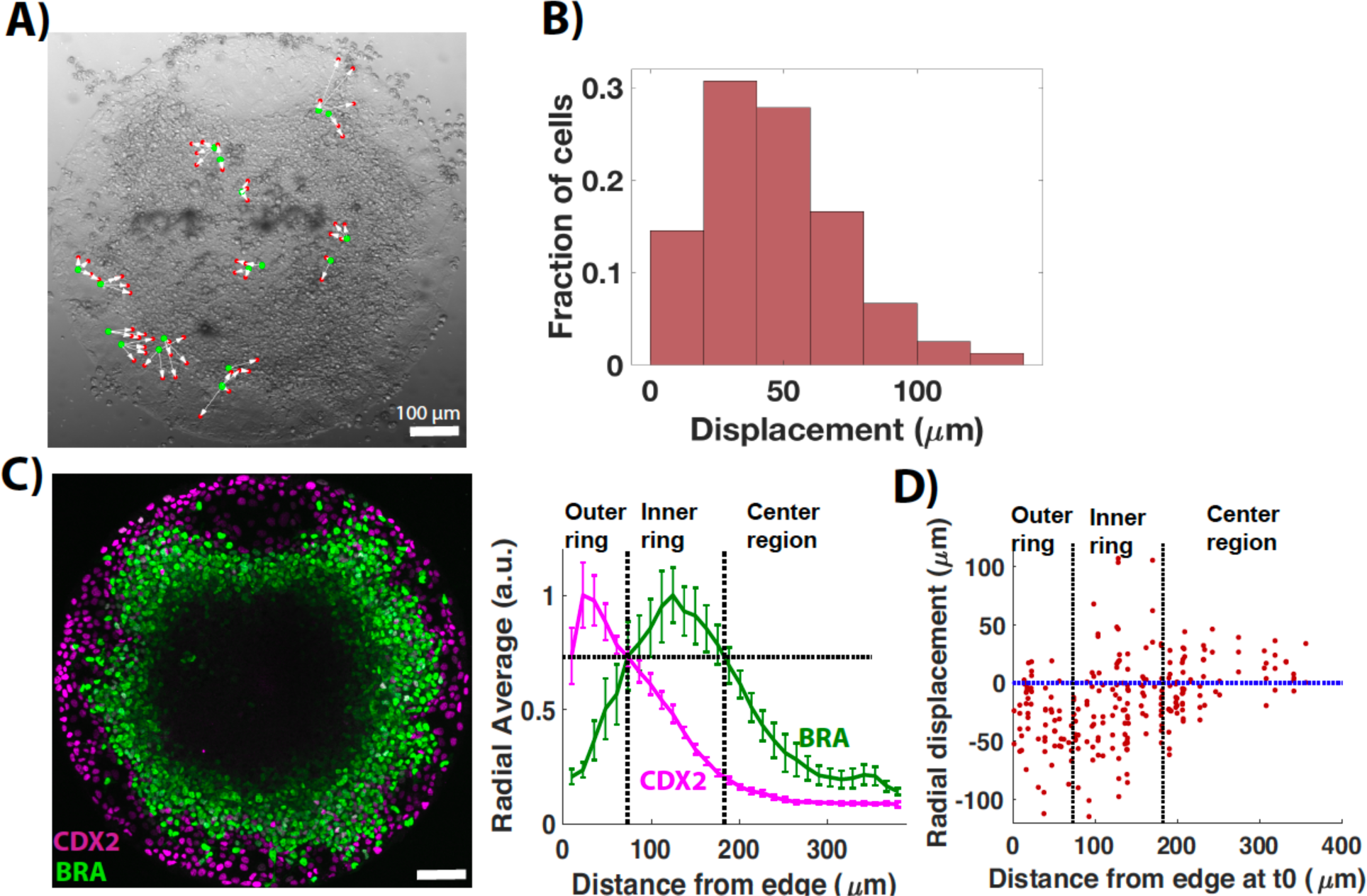
Inward movement of signaling activities is not due to movement of cells. **(A)** Snapshot from time-lapse imaging at 47h post BMP treatment in the bright field channel. Green and red dots represent initial and final positions respectively, of labeled cells that were correctly tracked throughout imaging. **(B)** Histogram of displacement of tracked cells. n = 84 cells. **(C)** Image of colony (shown in A) immunostained for CDX2, BRA post live cell imaging. Quantification represents average nuclear intensities of indicated markers normalized to DAPI as a function of radial position. Error bars represent standard error of the mean. N = 6. The x coordinate of the point of intersection of CDX2 and BRA curves (73.65 um) was chosen as the region bound for the high CDX2 region (outer ring, vertical dotted line passing through intersection point). The y coordinate of the intersection point was chosen as the threshold to define intensity level for high BRA region (horizontal dotted line). The x coordinates of the intersection points of threshold line and the Brachyury curve (73.65, 183um) were chosen as the region bounds for high BRA region (inner ring, vertical dotted lines). **(D)** Radial displacement of cells as a function of their starting position. Radial displacement = distance of the cell from the center at end of imaging - distance of the cell from the center at the start of imaging. The vertical dotted lines represent three different regions as defined in C. The horizontal dashed line separates the cells moving towards the edge from cells moving towards the center. See also S5, movie9

Quantifying cell movement revealed that the displacement of cells averages only about two cell diameters from the starting position (Figure 6B). Although cells in the outer region of the colony move inwards (Figure 6D, S5), the physical movement of cells occurs over a smaller distance than the movement of signaling domains, with both WNT and NODAL signaling domain moving at least 100µm inwards during this time (Figure 2, (15)), suggesting that inward movement of paracrine WNT and NODAL signaling activities is not a result of active cell migration. Besides, patterns of cell division are very similar in the three regions (Figure S5A, B), suggesting that differential cell growth does not play a role in fate patterning.

### Mathematical modeling reveals that signaling waves are not caused by Turing instability

Our results suggest that BMP treatment triggers waves of paracrine WNT and NODAL signaling activity that move towards the colony center, independently of cell movement. Both WNT and NODAL signaling are known to activate the production of their own ligands and inhibitors (29,30), and previous results have suggested that these inhibitors are essential for patterning (5,10,16). Previous work has also suggested that these activator-inhibitor motifs can function to render the state of homogenous signaling unstable driving pattern formation (31–35). Using mathematical modeling, we tested if such models could be a plausible mechanism behind the formation of signaling waves. For simplicity, we included only one activator-inhibitor pair, where the activator-inhibitor symbolizes a composite of WNT/NODAL and their secreted, diffusible inhibitors (Figure 7A). To avoid the effects of boundary conditions, we simulated the colony in a larger lattice with diffusion and protein degradation allowed throughout the lattice, and protein production occurring only within the colony. We started with initial conditions of high signaling at the colony border, which reflects both the initial higher activity of NODAL in this region (Figure 3A), and the initiation of both the WNT and NODAL fronts there (Figure 2, (15)).

**Figure 7:**
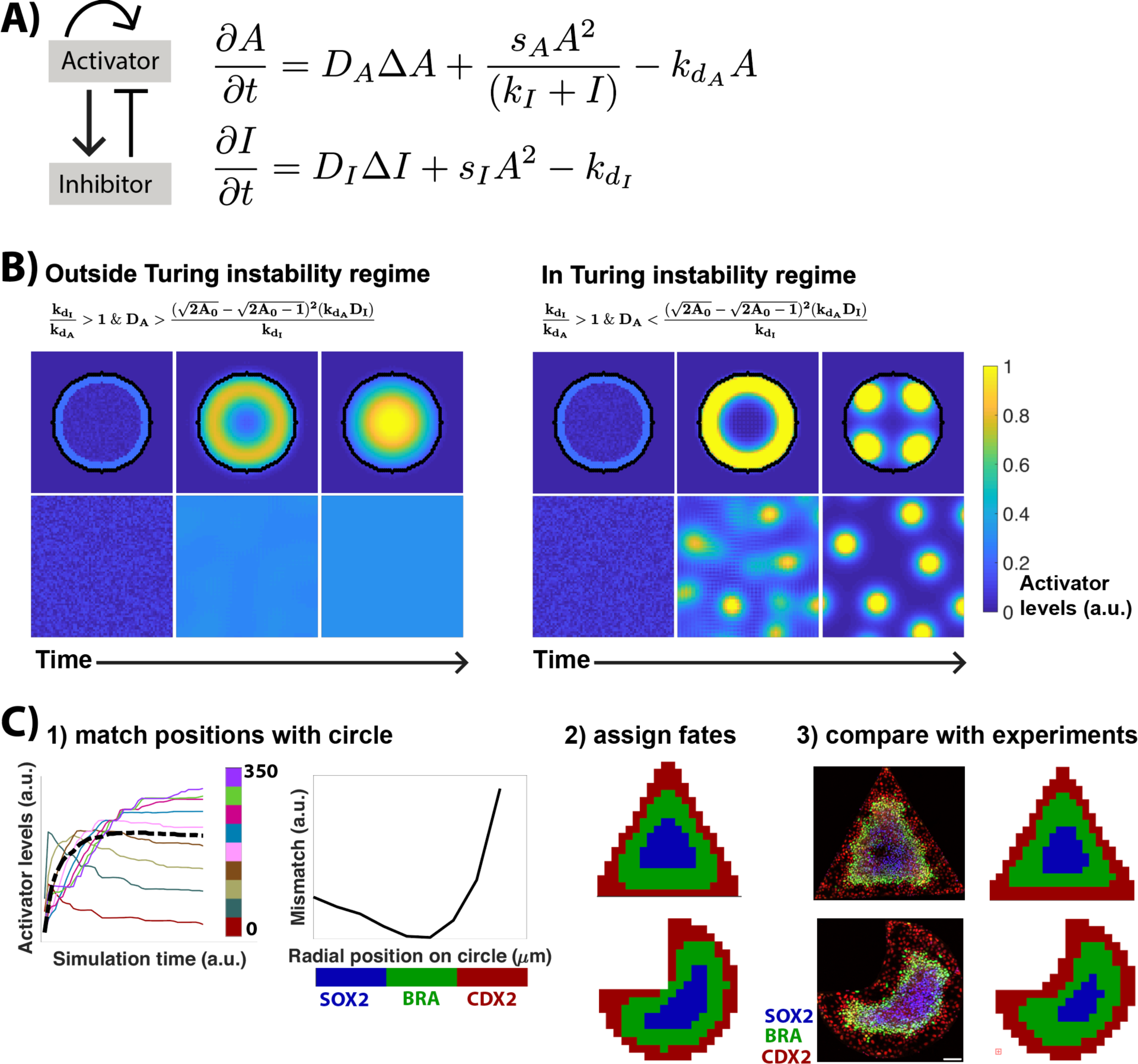
Mathematical modeling reveals that signaling waves are not caused by Turing instability. **(A)** Schematic and equations of the model. **(B)** Evolution of activator levels from the initial state to the steady state. Inequalities define parameter regimes that display each of the two behaviors (refer modeling supplement). Simulation parameters: s_A_= 0.01, k_I_ = 1, k_A_ = 0, kd_A_ = 0.001, D_I_ = 0.4, s_I_ = 0.01, kd_I_ = 0.008. (left) D_A_ = 0.014, (right) D_A_ = 0.0025 for outside and inside Turing instability regime respectively. Simulation domain: 190*190 pixels square lattice with periodic boundary conditions and random distribution of activator/inhibitor as initial conditions. To simulate the model in circular colonies, a circle (radius: 25 pixels) is defined at the center of the lattice. Assumptions: 1. Reactions take place only inside the circle; diffusion takes place in the entire lattice. 2. Activator/inhibitor degrade faster outside the circle (kd = 0.01). All images correspond to the center 90*90 pixels of the simulation lattice. **(C)** Workflow to assign fate territories on colonies of different shapes. 1. Simulated activator levels as a function of time for one position in the triangular colony (black dashed line) and for different radial positions in the circular colony (colored curves). Colorbar represents radial position (distance from the edge(µm)). Mismatch is calculated as the sum of squared differences between activator evolution in time for the position in the triangular colony and different radial positions in the circular colony. 2. The position in the triangular colony is assigned the fate of the most similar position in the circular colony. The procedure is used to assign fates to all positions in triangular and pacman colonies. 3. hESCs immunostained for CDX2, BRA, SOX2, after 44h of BMP treatment on triangular and pacman colonies. Immunostaining data from n=18 colonies was used to calculate average fate territory maps shown adjacent to image. Scale bar = 100µm. See also S6, movie10,11,12,13.

The simple reaction-diffusion model recapitulates the experimentally observed inward movement of signaling activities in a broad parameter range (Figure 7B, Model supplement, Movie S10-S13). To our surprise, the range in which the model agrees with the data lies entirely outside the regime with a diffusion-driven instability (Figure 7B). That is, the models that generate waves of signaling activity, when simulated on a lattice with periodic boundary conditions, do not generate patterns. Conversely, in the regime where patterns form as the result of a Turing instability, signaling waves are not seen on restricted geometries, thus indicating that signaling waves are not a result of a Turing instability. We confirmed these results analytically by deriving conditions for the stability of the homogenous state (Figure 7B, Model supplement). We also tested a stripe-forming Turing model, which, like the spot-forming model, does not recapitulate the signaling waves in the circular colony (Figure S6A), further corroborating that signaling waves are not a result of Turing instability. Instead, the waves result from an initial activation at the colony boundary, and the system moves towards a steady state of WNT/NODAL signaling that is homogenous. The differences between the edge and center of the colony at steady state result only from diffusive loss of signals at the colony boundary. Thus, in the experimental regime, the homogeneous steady state is stable, and initiation of signaling at the colony boundary leads to an expanding wave-like behavior of signaling activities as the system tends towards this homogenous state. Spatial patterns in signaling activity at steady state result only from boundary effects and do not correlate with the cell fate pattern.

As a test of the model, we simulated fate patterns on colonies with different geometries. Simulations of different shapes can be performed without changing the model, and only require adjusting the boundary conditions to reflect the new shape. Thus, there are no free parameters involved in this test of the model. To define fate patterns in the new shape, we matched each position in the new shape with the position in a circular micropatterns that had the most similar simulated signaling dynamics and assigned it the fate associated with that position (Figure 7C, S6).

The predicted fate patterns matched the experimentally observed ones showing an inward expansion of cell fates at the corners of the colonies in both cases (Figure 7C). Thus, our model can predict signaling dynamics and cell fate patterns on varying geometries with no further input.

## Discussion

Spatially confined hESCs treated with BMP4 self-organize to form radial rings of distinct germ layers: an outer ring of extra-embryonic cells, followed consecutively by rings of endoderm, mesoderm and ectoderm cells, thus, recapitulating the patterning associated with gastrulation *in vitro* (5). Recent studies have shown that edge restricted BMP signaling drives extra-embryonic differentiation (16,17). Here, we show that continuous BMP signaling at the colony edge initiates a wave of WNT signaling that moves towards the center of the colony. WNT signaling in turn initiates a NODAL wave, which, once initiated, propagates independently of both BMP and WNT signaling. We further show that the duration of BMP signaling specifies the position of mesodermal differentiation and continuous signaling through both the WNT and NODAL pathways synergizes to achieve maximal differentiation.

Although WNT and NODAL signaling waves are necessary for mesodermal differentiation, both waves move further towards the center of the colony than the ring of mesodermal differentiation (Figure S1C, 2B,3C,4B), suggesting that mesodermal differentiation does not dependent on a particular threshold of signaling levels. What, then, defines the boundaries of mesodermal ring? Given that both WNT and NODAL fronts move inwards at different speeds, with NODAL moving faster than WNT ((Heemskerk et al., 2017), Figure 2), one hypothesis is that the interval between the WNT and NODAL fronts reaching a cell is a key parameter in determining mesoderm differentiation, consistent with a recent study on NODAL dynamics (36). Another hypothesis is that since BMP signaling is activated homogeneously before being edge restricted to the edge (15), it is possible that the timing between BMP signaling and the WNT and NODAL waves determines the fate positions. Both hypotheses suggest that it is the timing of signaling, and in particular the relative timing of different pathways, rather than the signaling levels themselves that determine the fate patterns. Notably, as discussed below, our data are not consistent with WNT or NODAL forming Turing patterns as they eventually evolve to a homogenous state that encompasses the entire colony (Figure 3A).

Reaction-diffusion based activator inhibitor models are theoretically capable of forming a wide range of patterns by a diffusion-driven instability that renders the homogeneous steady state unstable and transitions the system to a new steady state with spatial patterns (34,35). Homologues of NODAL and its inhibitor Lefty, when overexpressed in zebrafish embryo, have diffusion and degradation rates which are consistent with a Turing system (33). However, whether the NODAL and Lefty behave as a Turing system at endogenous levels remains unknown. Here, we show that both NODAL and WNT signaling activities settle to a homogeneous steady state throughout the gastruloid shortly after patterning the mesoderm, with the only regions of low activity being at the colony edge. Mathematical modeling suggests that temporally expanding activity is the result of initial activation at the colony edge combined with autoactivation, and that the low levels at the colony border at steady state arise from diffusive loss of WNT and NODAL ligands across the colony boundary. Neither feature reflects a diffusion-driven Turing instability. Previous studies have suggested that BMP and its inhibitor Noggin, which restricts high BMP signaling to the colony edges, function as a Turing system (6,16). Our results are potentially consistent with this idea, as active BMP signaling does reach a fixed length scale of activity, however, our results are also consistent with diffusive loss of Noggin from the edges combining with constant levels of exogenous BMP to produce this pattern of signaling. Differentiating between these models requires determining whether the BMP-Noggin system can break symmetry and initiate patterning in the absence of a colony border. In any event, once the BMP gradient is established, the mesendoderm is patterned by waves of paracrine WNT and NODAL signaling, not cells reading out a BMP gradient as was proposed previously (6). Thus, signaling dynamics, and not a stable spatial pattern of signaling activity driven by a Turing instability, control the self-organized patterning.

Our study also highlights a previously unappreciated role of WNT signaling in the differentiation of extraembryonic CDX2+ cells. We show that continuous WNT signaling is necessary for these cells to form in the outer ring of the colony. This is striking because WNT signaling never peaks in the colony edges. This suggests that either low levels of WNT signaling are instructive or WNT signaling affects extra-embryonic differentiation through secondary effects in a non-cell autonomous manner. Another possibility is the involvement of non-canonical WNT signaling, independent of β-catenin, in extra-embryonic differentiation, and future studies will be needed to address this.

The involvement of WNT signaling in CDX2+ differentiation also raises a question on the identity of these cells. One possibility is that these cells represent extra-embryonic mesoderm cells which originate in the proximal posterior of the mouse embryo where WNT signaling is high (1). But, unlike mesodermal cells, these CDX2+ cells do not pass through a BRA+ state as BRA+ cells do not start at the colony edge (15) and there is no large-scale outward movement of cells from the BRA+ ring (Figure 6). This suggests that the CDX2+ cells may represent a subset of extra-embryonic mesodermal cells that do not pass through a BRA+ state. Another possibility is that the CDX2+ cells represent trophectodermal cells. WNT signaling, however, is dispensable for mouse trophectoderm development (24) and also for *in vitro* trophectoderm differentiation of hESCs (25). This suggests that either *in vivo* human trophoectoderm development has different signaling requirements than the mouse, or that the requirement for WNT signaling is a tissue culture artifact. Future studies will be needed to determine the exact developmental status of these CDX2+ cells.

We observe limited cell movement as the gastruloids are patterned, which indicates that the signaling waves that instruct primitive streak differentiation are not a result of collective cell migration. This is in contrast to developing chick embryos where large-scale collective cell migration is integral to primitive streak formation (37). The evidence from the developing mouse embryo suggests an absence of collective migration immediately preceding primitive streak formation (38), and may be consistent with our results here. Absence of cell movement implies that signal wave result form either WNT and NODAL ligands moving through colonies, or that cells responding to signaling by producing ligands that activate their neighbors in a relay-like manner.

In a recent study, it was shown that WNT treated micropatterned hESCs also exhibit an inward expansion of WNT signaling activity. There, it was shown that a prepattern in which E-cadherin is low at the colony edge allows a high response to WNT signaling in edge cells. WNT combines with NODAL signaling to initiate an EMT, which downregulates E-cadherin in the neighboring cells and allows them to respond to the exogenous WNT. Thus, when microcolonies are stimulated with WNT, a wave of EMT propagates inwards and underlies the apparent movement of the WNT signal in the colony (39). In contrast, here we show that in BMP treated micropatterned hESCs, the inward expansion of WNT signaling activity does not start at the colony edge (Figure 2) where E-cadherin would be low. Further, the expansion of WNT signaling continues in the absence of an EMT or paracrine NODAL signaling (Figure 4F,G IWP2 @15h), suggesting that a similar E-cadherin mediated mechanism is unlikely to be involved. As the inward movement of WNT signaling activity continues even after inhibition of WNT secretion (Figure 4A-E IWP2@15h), it is possible that the wave of WNT signaling results from the inward movement of WNT produced at the colony edges prior to secretion inhibition. Inhibiting WNT secretion prevents autoactivation of the pathway as these ligands move inward, thus lowering the magnitude of the response, but leaving the range of signaling unaffected (Figure 4B-E). If true, this would indicate that WNT proteins have the potential to activate signaling at long-range, as recently observed in the *C. elegans* embryo (40), and in contrast to the short range activation in the intestinal crypt (41). The linear scaling in time of the movement of WNT signaling in the colony, even in the absence of secretion of new WNT ligands, suggests that WNT ligands do not move by passive diffusion, as this would be expected to scale with the square root of time. Future studies will be needed to dissect the mechanisms underlying the movement of both WNT and NODAL signaling activities.

Although recent technical advances have made it possible to image post implantation mouse embryos at cellular resolution (42), ethical challenges make it impossible to perform similar studies in a developing human embryo (43,44). Thus, gastruloids offer a unique tool to investigate human gastrulation. Using this tool, we have shown that WNT and NODAL, two signaling pathways that are integral to primitive streak formation, have similar spatial dynamics but distinct temporal regulation by upstream signaling. We further show that neither WNT nor NODAL forms a spatial pattern in signaling activity that directly maps to the resulting fate patterning. Instead, our data suggest that fate patterning results from the dynamics of signaling activities. Thus, our study reveals that signaling dynamics in the absence of a diffusion driven Turing instability has the potential to drive self-organized fate patterning. In the vertebrate neural tube, mutually inhibitory interactions in the gene regulatory network that decodes Sonic hedgehog signaling are integral to establishing spatial fate patterns (45). It will be interesting to decipher the gene regulatory network that decodes position and hence final fates from the dynamics of signaling events during the self-organized fate patterning of human gastruloids.

## Acknowledgements

We thank Idse Heemskerk for helpful discussions on the project, Eric Siggia for early discussions and versions of the simulation code, Elena Camacho Aguilar, Eleni Anastasia Rizou, and Joseph Massey for careful reading of the manuscript, Cecilia Guerra for helping with experiments, and all the members of Warmflash lab for helpful feedback. This work was funded by Rice University and grants to AW from CPRIT (RR140073), NSF (MCB-1553228), NIH (R01GM126122), and Simons Foundation (511079).

## Author Contributions

Conceptualization: A.W and S.C; Methodology: S.C, L.L and A.W.; Software: S.C, R.G, A.W; Investigation: S.C, L.L; Writing – Original Draft, Review and Editing: S.C and A.W. Funding acquisition, Project Administration, Supervision: A.W.

## Materials and Methods

### Experimental System

#### Cell lines

Experiments were performed using ESI017 (obtained from ESI BIO, RRID: CVCL_B854, XX) human embryonic stem cell (hESC) line. For WNT signaling dynamics, the ESI017 GFP-β-catenin cell line as described in (26) was used. For cell tracking experiments, transgenic RUES2 cell line (RUES VENUS:H2B) (a gift from from Ali Brivanlou, Rockefeller, RRID: CVCL_B810, XX) was used.

### Methods

#### Routine cell culture

All cells were grown in the chemically defined medium mTeSR1 in tissue culture dishes and kept at 37°C, 5% CO_2_ as described in (17). Cells were routinely passaged and checked for mycoplasma contamination also as described in (17).

#### Micropatterning

Micropatterning experiments were performed on either micropatterned chips or 96 well micropatterned plates obtained from CYTOO (shapes, 96well plate, circles). In both cases, hESCs were seeded onto micropatterned surfaces coated with 5µg/ml Laminin-521 using the mTeSR1 protocol described in (19). Following seeding, cells were either treated with 50ng/ml BMP4 (gastrulation assay, control sample) and/or with reagents as described in the text.

#### Immunostaining

Immunostaining followed standard protocols as described in (17). Primary and secondary antibody were diluted in the blocking solution as described in (5,17). Dilutions are listed in the reagents table).

#### Creation of NODAL knockout (NODAL-/-) cells

We introduced a mutation in exon 1 of the NODAL gene in ESI017 hESCs using CRISPR-Cas9. A guide RNA(sgRNA) directed towards NODAL exon1 was designed using benchling. The single strand oligonucleotides for sgRNA were NODAL_sgRNA_Forward:-CACCGGGCCCACCAGGCGTGCAGA; NODAL_sgRNA_Reverse:-AAACTCTGCACGCCTGGTGGGCCC These oligonucleotides were annealed and inserted into the PX459 vector through BbsI restriction sites using standard restriction and ligation protocols. The insertion was verified using DNA sequencing. DNA was nucleofected into 8*10^5^ hESCs using P3 Primary Cell 4D-Nucleofector^®^ X Kit L (Lonza). Following nucleofection, cells were transferred into mTeSR1 with 10µM ROCK inhibitor. Cells were selected by adding 1µg/ml Puromycin to the nucleofected cells the subsequent day. To increase survival of selected cells, mTeSR1 was supplemented with CloneR after one day of antibiotic selection. After five-six days in mTesr1 and CloneR, single colonies were picked and transferred to a 24 well plate. Genomic DNA from selected cells was extracted using DNeasy Blood and Tissue Kit. The genomic region around the NODAL gene was PCR amplified using primers: Forward primer: 5’-TTGCAGCCTGAGTGGAGAGG-3’; Reverse primer: 5’-AACCCACAGCACTTCCCGAGTC-3’ The PCR product was cloned using the Invitrogen™ TOPO™ TA Cloning™ Kit. The PCR product and the individual colonies from TOPO cloning sent for DNA sequencing. Sequencing results showed the presence of only two distinct mutations (Figures S2) on the NODAL genomic locus, suggesting the enrichment of a monoclone with a distinct mutation on each allele. The absence of functional NODAL protein was verified by Western blot.

#### Western blot

Wild type ESI017 hESCs and NODAL-/- hESCs were treated with 10uM WNT agonist CHIR99021(46) for 20h, which activates WNT-β catenin pathway and the transcription of downstream target genes like NODAL (28). After treatment, cells were washed with PBS and lysed with cOmplete Lysis-M solution. Following lysis, cells were mixed with 2x Laemmli Sample Buffer supplemented with 200mM Dithiothreitol. The samples were denatured by heat at 95°C for 5 minutes, and loaded to 4–20% Mini-PROTEAN^®^ TGX Precast Gel (Bio-Rad). After electrophoresis at 120 V for 90 to 120 minutes, the sample was transferred to polyvinyl difluoride (PVDF) membrane. The PVDF membranes were blocked in PBST with 5% non-fat milk. The primary antibodies (NODAL and beta actin) were dissolved in PBST with 2% non-fat milk and incubated on the membrane at 4°C overnight. The membrane was washed 3 times with DPBST (1× DPBS with 0.1% Tween 20). Horseradish peroxidase conjugated secondary antibodies were applied to the membrane and incubated at room temperature for 1 hour. The membranes were washed 3 times with PBST and the signal was detected by using ECL Western-Blotting substrate and Amersham Hyperfilm ECL.

#### Imaging

*Live cell imaging:* For cell tracking experiment, cells with a nuclear fluorescent marker (RUES2-VENUS-H2B) were mixed with unlabeled cells (ESI017) in the ratio 1:100, seeded onto to a micropatterned chip kept in a holder (CYTOO) and treated with 50ng/ml BMP4 as described in (19). Six colonies of 800µm diameter were imaged with a 10X, NA 0.40 objective on an Olympus laser scanning confocal microscope. 5-8 z-slices were acquired for each position every 10 minutes from 20h to 47.5h post BMP4 treatment. For determining WNT signaling dynamics during differentiation, GFP-β-catenin-hESCs were seeded and treated as above. Nine colonies of 800µm diameter were imaged with a 20X, NA 0.75 objective on a laser scanning confocal microscope with 5 z-slices acquired per position every hour from 3h to 47h post BMP4 treatment (Figure 2). For comparing WNT signaling dynamics across multiple conditions, GFP-β-catenin-hESCs were seeded in multiple wells of a 96 well micropatterned plate (CYTOO), with one experimental condition per well. Colonies of 700µm were imaged at 20X resolution, NA 0.75 on a spinning disk confocal microscope, with 5 z slices acquired per position every 30 minutes from 3h to 47h post BMP4 treatment. Cells were treated with 200nM LDN or 5µM IWP2 at the indicated times (Figure3, 4). The number of colonies imaged for each condition were as follows: LDN@0h:- 3, LDN@11h:- 3, LDN@23h:- 5, Control with no LDN:-13; IWP2@15h:- 5, IWP2@30h:- 5, Control with no IWP2:- 5, where time in each condition represents time post BMP4 treatment when the indicated reagent was added. For all the above-mentioned experiments, cells were maintained at 37C and 5% CO_2_ during imaging.

*Fixed cell imaging:* All the immunostaining data was acquired by imaging entire fixed micropatterned chips and 96 well micropatterned plates using tiled acquisition with a 10X, NA 0.40 objective on an Olympus IX83 inverted epifluorescence microscope. For data visualization purposes, sample images for each condition were acquired using 20X resolution NA 0.75 on laser scanning confocal microscope. Raw images for immunostaining data in each main figure correspond to images taken at 20X and average plots represent quantification of images taken at 10X.

### Quantification and Analyses

All experiments were performed at least twice with consistent results. The data and analyses in each figure belong to one experiment. Sample size was not pre-determined and no statistical tests were used to determine significance of results. Circular colonies with a non-radial cell density pattern at the end of 44h of BMP4 treatment were excluded from analyses. The number of colonies included in each analysis (N) is mentioned in the figure legends. For images taken at 20X magnification with multiple z-slices; background subtraction, maximum z projection and alignment were performed as described in (5). Colony images obtained after alignment were analyzed as described in Figure S7 using custom made MATLAB scripts.

For the cell tracking experiment, cells were tracked using the tracking workflow in Ilastik 1.2.0 (47). Of the 165 labeled cells in six colonies, at least one daughter cell of 84 cells was tracked correctly during the entire course of imaging. Cells that died, went out of focus, or were mis-tracked were excluded from analyses.

### Data and Software

All the raw data is available upon request. MATLAB scripts for analyzing experimental data can be obtained from https://github.com/sc65/CellTracker and for running simulations from https://github.com/sc65/pde_simulation.

### Resources Table

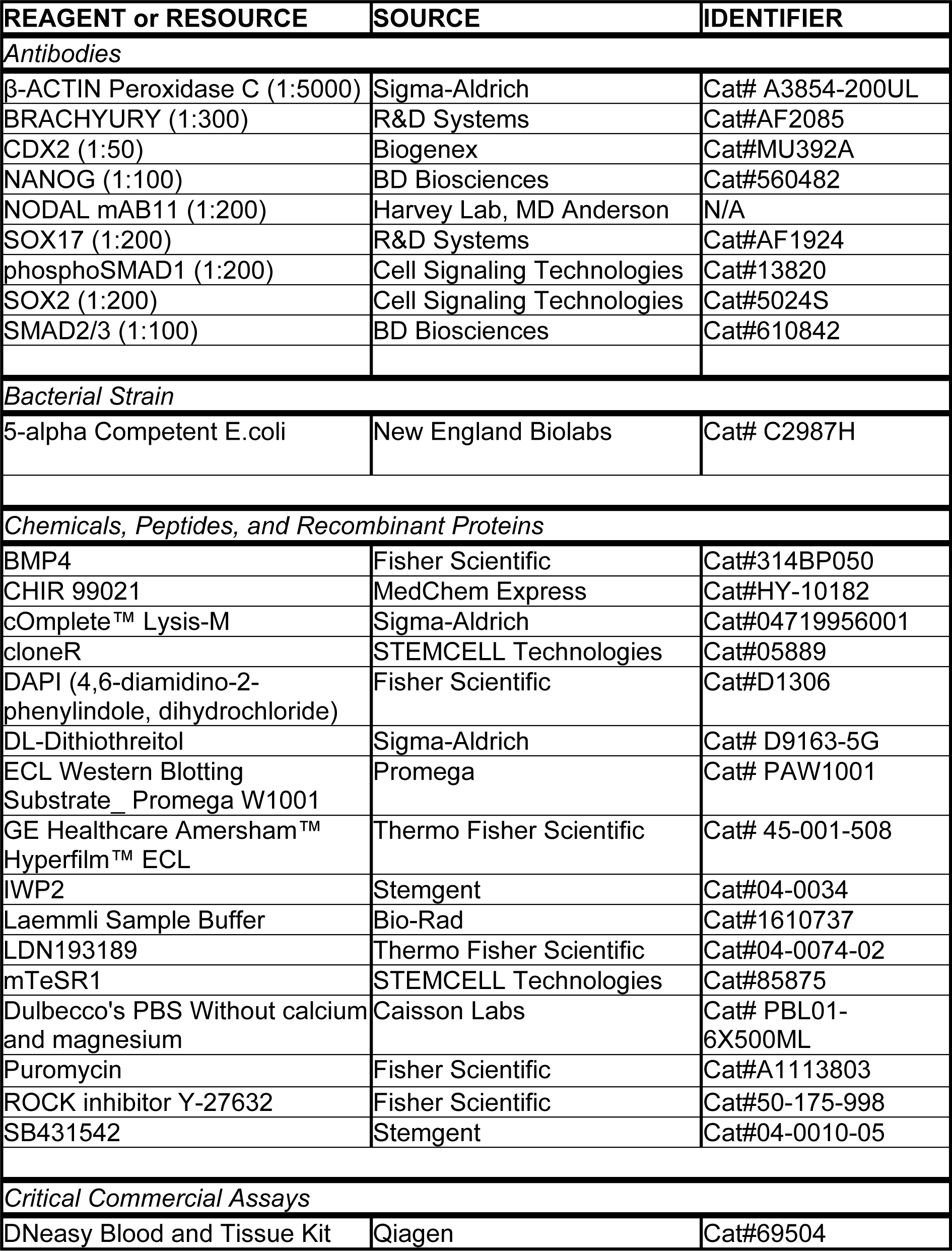
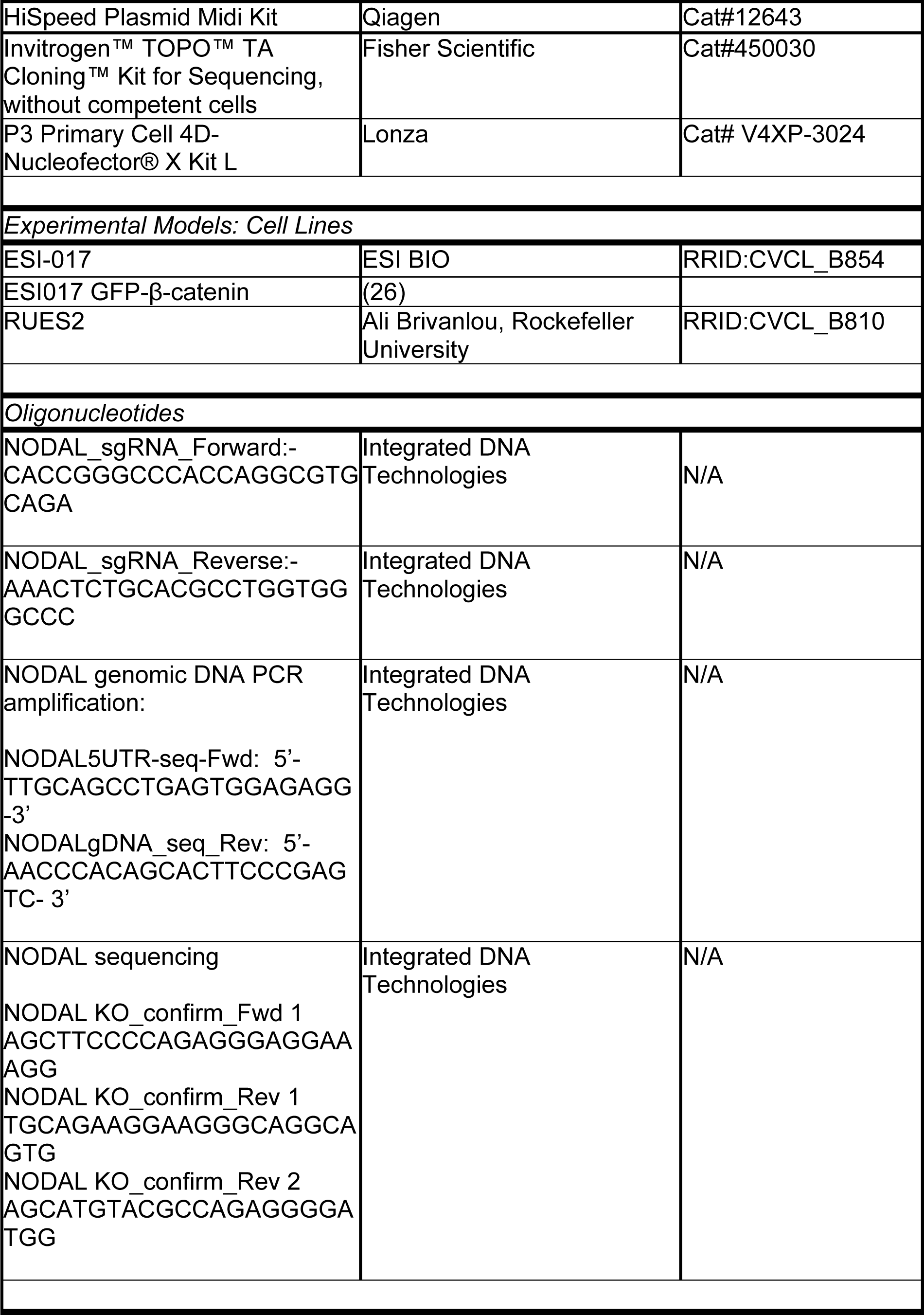
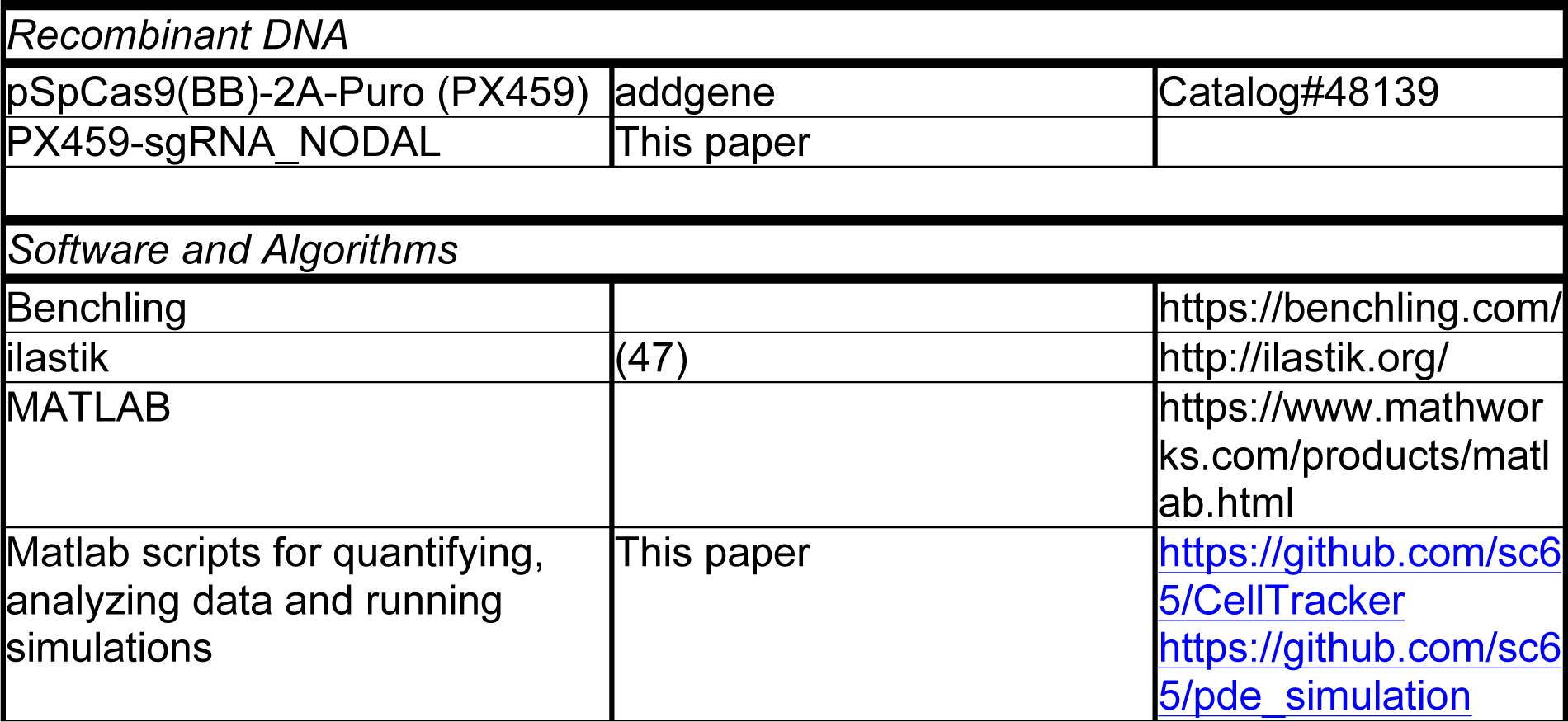

