## Supplementary material for "Dissecting the dynamics of signaling events in the BMP, WNT, and NODAL cascade during self-organized fate patterning in human gastruloids": Figures S1-S7 and supplemental text

### Supplementary figures

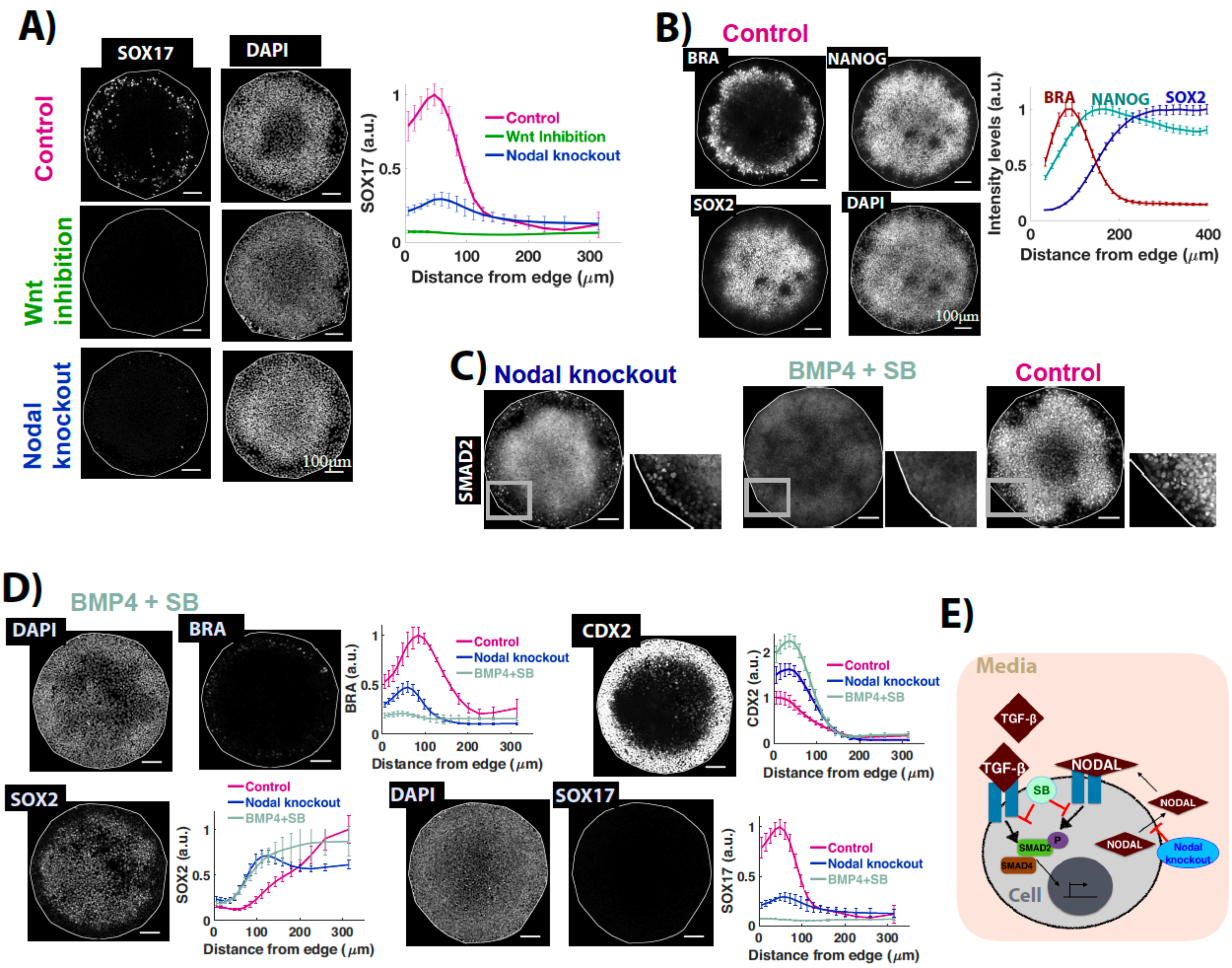


**FigureS1: Fate patterning requires Wnt and Nodal signaling, related to figure1**
**(A)** Images of samples immunostained for SOX17 in different conditions-Control and Nodal$^{-/-}$ cells were treated with 50ng/ml BMP4. Wnt inhibition indicates treatment of wildtype cells with 50ng/ml BMP and 5 μM IWP2. All samples were fixed 44h post treatment. Quantification represents intensity levels of indicated markers normalized to DAPI, averaged at different positions along the colony radii (radial averages, S7). Error bars represent standard error of the mean. N >= 10. Colony diameter= 700μm. **(B)** Images of the samples immunostained for NANOG, SOX2 and BRA after 44h of BMP treatment. Quantification represents intensity levels of indicated markers normalized to DAPI, averaged at different positions along the colony radii. N = 24. **(C)** Images of samples immunostained for SMAD2 after 44h of BMP4 treatment in different conditions as indicated above colonies and defined in D. Region marked by the square is zoomed in small images adjacent to each condition. Scale bar = 100μm. **(D)** Images of samples immunostained for indicated markers after 44h of treatment with 50ng/ml BMP4 and 10μM SB (SB treated). Quantification represents intensity levels of indicated markers normalized to DAPI, averaged at different positions along the colony radii in the SB treated, Control and Nodal knockout samples. N>=10. **(E)** Schematic showing Nodal, TGF-b binding to the receptors and nuclear translocation of signal transducer SMAD2. TGF-beta inhibitor SB, blocks the activity of TGF-beta type 1 receptors like ALK4,5,7 and thereby inhibits downstream signaling. Nodal knockout cells are incapable of Nodal secretion.

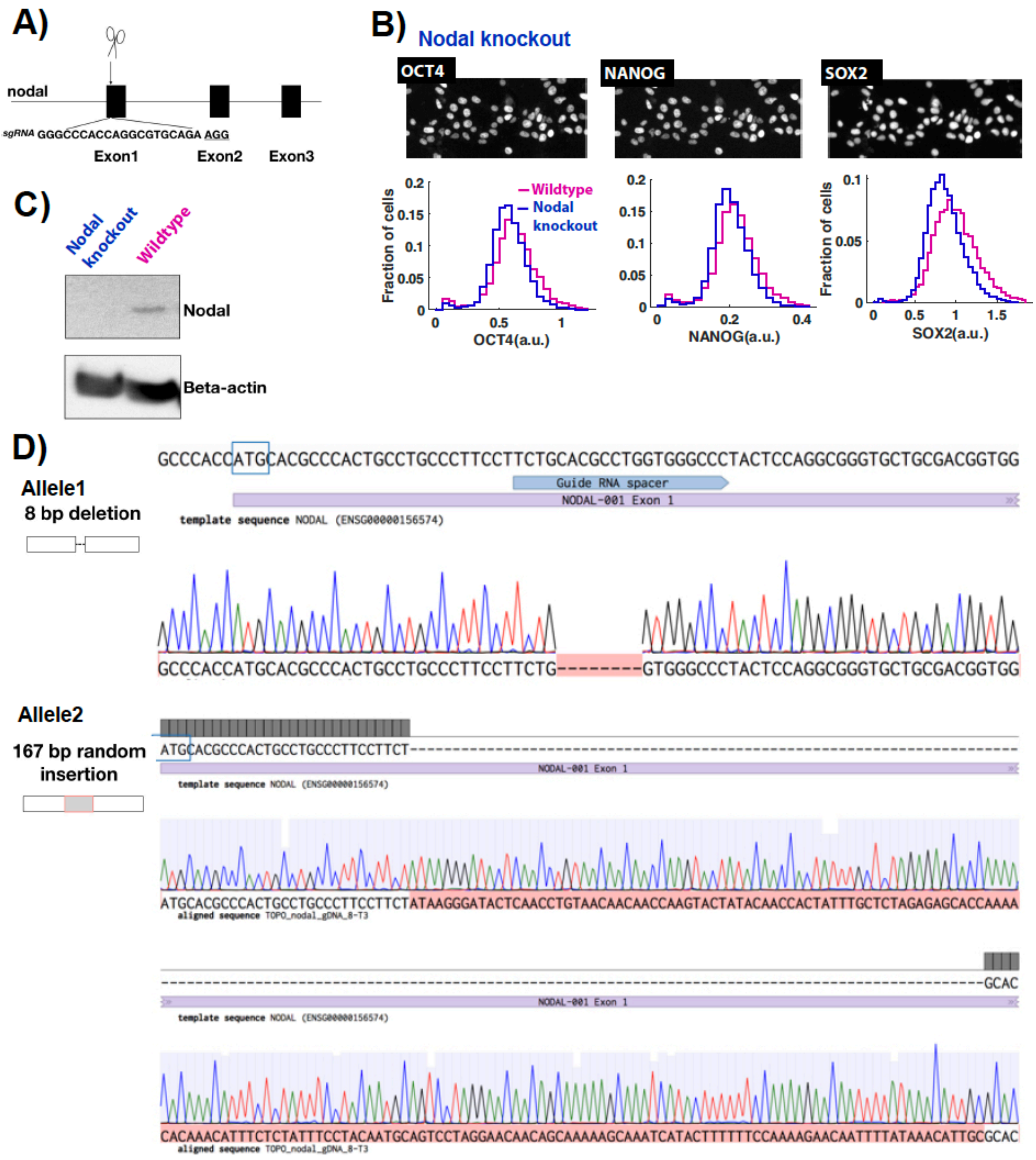


**Figure S2: Creation and Validation of Nodal knockout cells, related to figure 1**
**(A)** sgRNA used to make a double stranded break on exon1 of endogenous Nodal gene. **(B)** Images of Nodal knockout cells immunostained for pluripotency markers OCT4, NANOG, SOX2. Histograms represent marker levels normalized to DAPI. N >1000 **(C)** Western blot for Nodal following treatment with 10μM CHIR in Wildtype ESI017 cells and Nodal knockout cells. **(D)** Genomic sequence of Nodal locus in Nodal knockout cells.

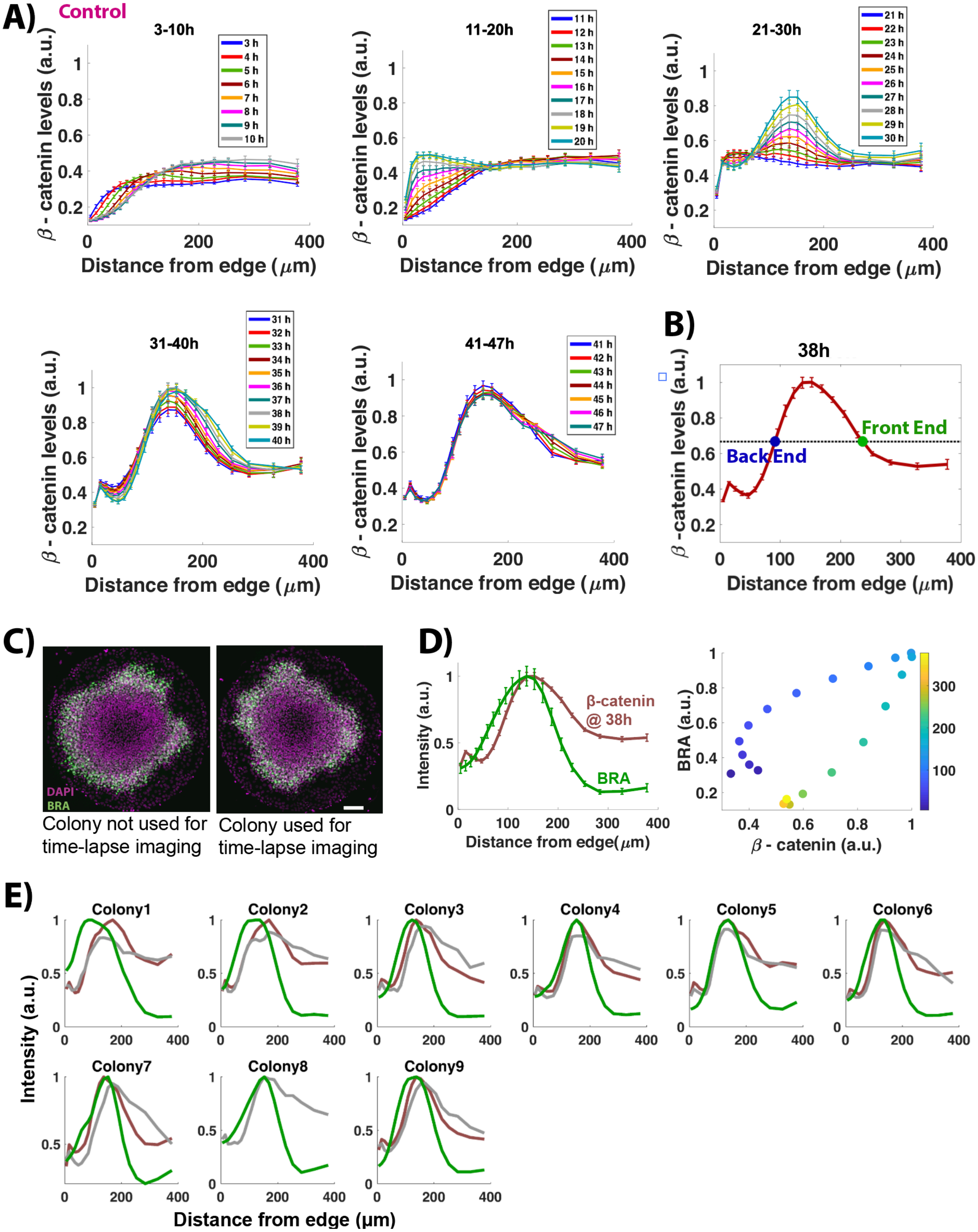
A)
Control
3-10h
11-20h
21-30h
31-40h
41-47h
β - catenin levels (a.u.)
Distance from edge (μm)
B)
38h
Front End
Back End
C)
DAPI
BRA
Colony not used for time-lapse imaging
Colony used for time-lapse imaging
D)
Intensity (a.u.)
β-catenin @ 38h
BRA
Distance from edge(μm)
BRA (a.u.)
β - catenin (a.u.)
E)
Colony1
Colony2
Colony3
Colony4
Colony5
Colony6
Colony7
Colony8
Colony9
Intensity (a.u.)
Distance from edge (μm)

**Figure S3; Wnt signaling dynamics in the control sample (47h BMP4 treatment), related to figure2**
**(A)** Average non-membrane beta-catenin levels as a function of radial position at different times post BMP treatment. **(B)** Threshold signaling (dotted line) defined as the half max of average non-membrane beta-catenin levels at time point when signaling peak is the highest (38h). n = 9. Error bars indicate standard error. (**C**) Images of colonies immunostained for BRA and DAPI. Scale bar: 100μm. **(D)** (Left) Average BRA intensity levels and non-membrane β-catenin at highest signaling as a function of radial position. (Right) Average BRA intensity levels as a function of non-membrane β-catenin color-coded by edge distance (μm). **(E)** Average BRA intensity levels, non-membrane β-catenin at highest signaling (red curve) and last time point (47h, gray curve) as a function of radial position for individual colonies. In Colony 8, highest signaling occurs at the last time point.

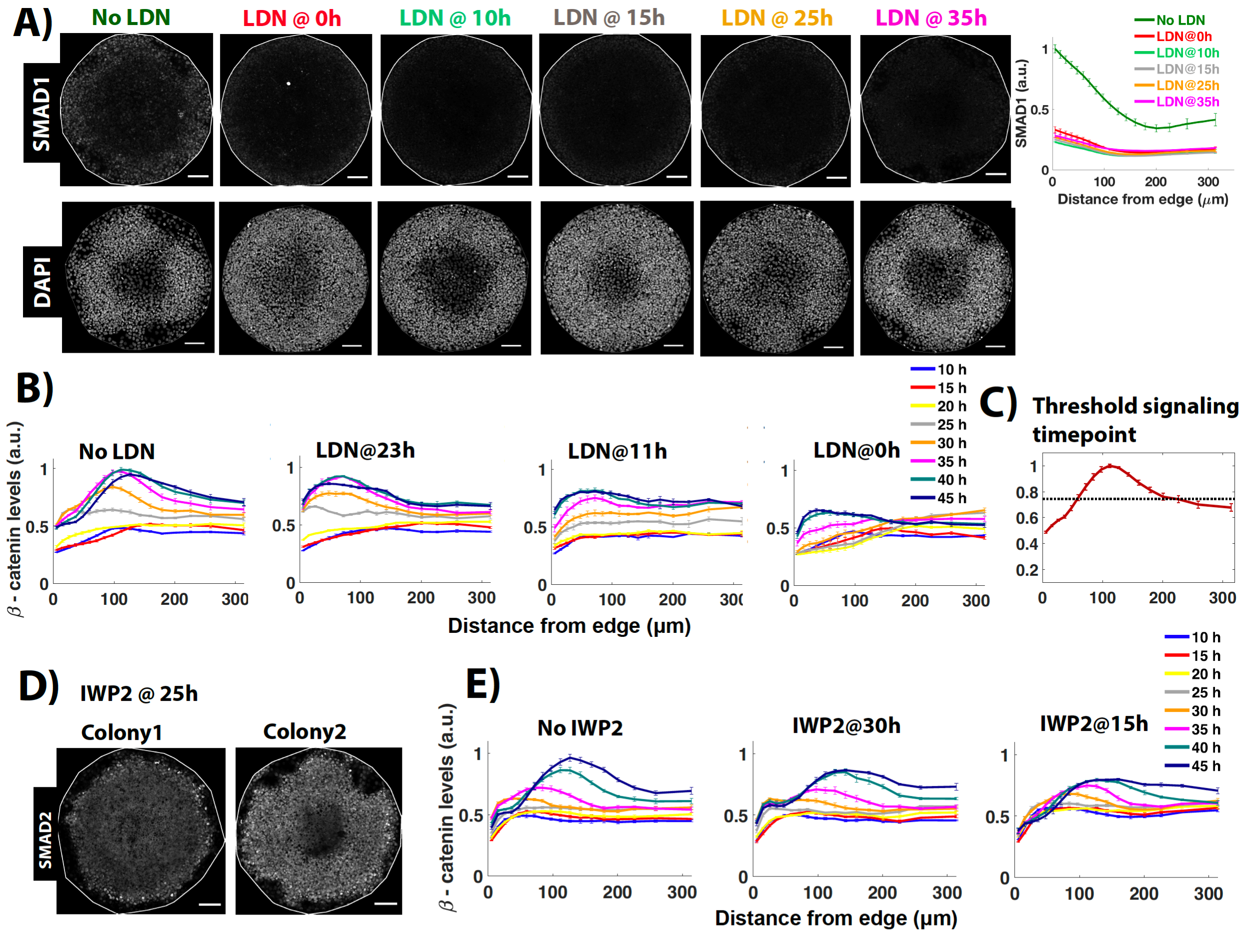


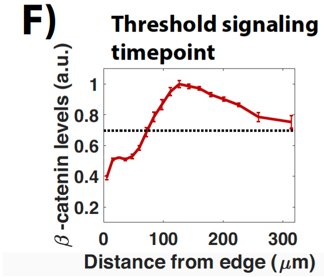


**Figure S4: Wnt signaling dynamics in LDN and IWP2 treated samples related to figure3,4**

(**A**) Images of colonies immunostained for SMAD1 and DAPI after 44h of BMP treatment. The time between BMP4 and LDN addition is indicated above the image. No LDN was added in the control sample. Quantification represents average nuclear intensities of indicated markers normalized to DAPI as a function of radial position. N>=5. **(B,E)** Average non-membrane β-catenin levels as a function of radial position. The time in the legend represents time post BMP treatment being analyzed in each curve. The time above the curves indicate the time between BMP4 and LDN/IWP2 treatment. No LDN/IWP2 was added in control. **(C,F)** Average non-membrane β-catenin levels at the time point when signaling is highest in the control sample. This time point was used to define threshold signaling levels (dashed line) for each condition. **(D)** Images of colonies immunostained for SMAD2 at 44h post BMP treatment. IWP2 was added at 25h post BMP treatment. For all radial averages plot, error bars represent standard error. Scale bar = 100um.

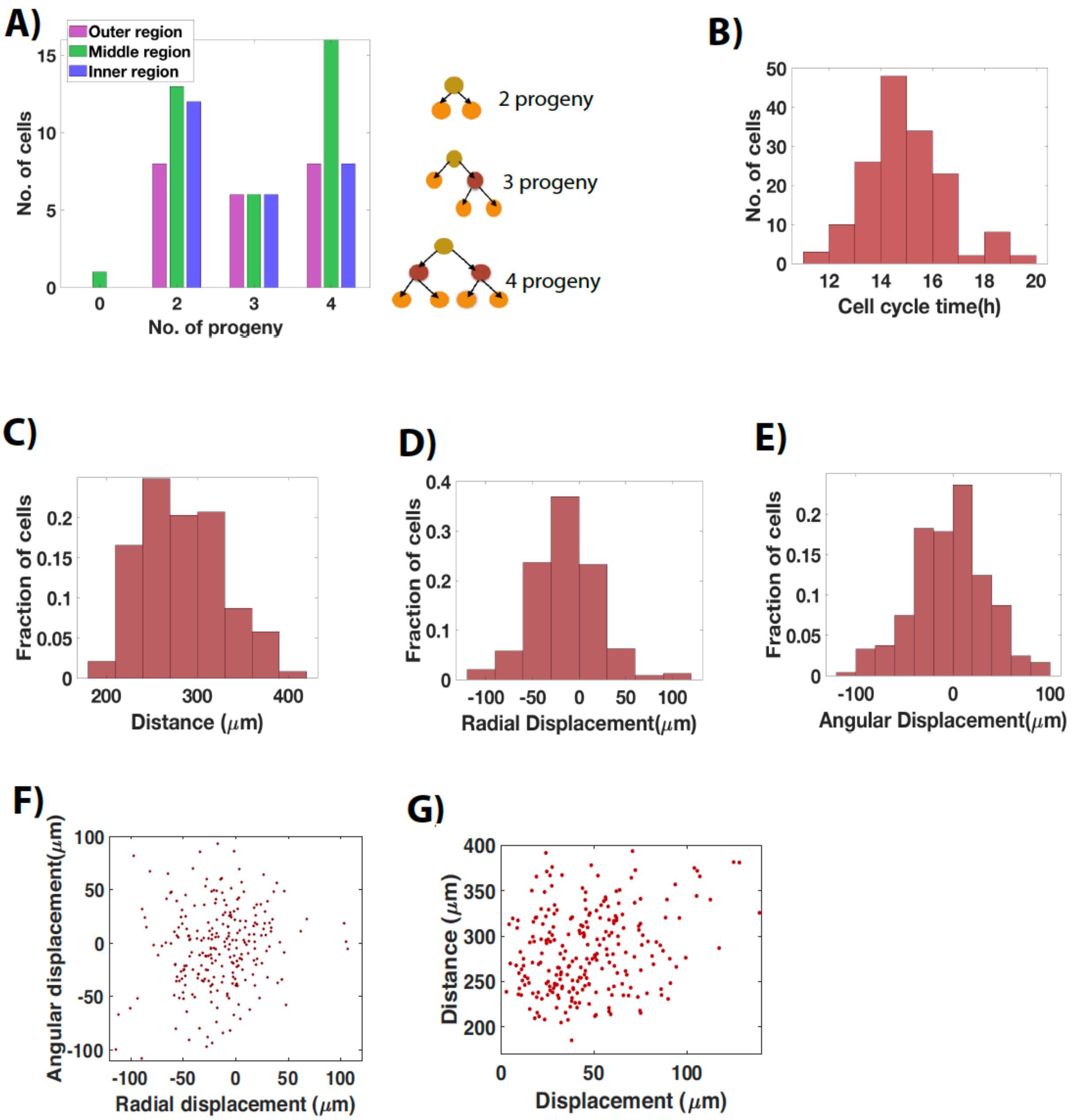


**Figure S5: Cell division and cell movement during fate patterning, related to figure6**
**(A)** Number of progeny of tracked cells that start in the outer, inner or center regions as defined previously (Figure6). No significant difference between cell division trends across three regions. kstest2 returned 0 for all three comparisons. 0 progeny: No cell division, 2 progeny: 1 cell division, 3 progeny: 1 daughter cell divides, 4 progeny: both daughter cells divide (pictorial representation adjacent to figure). **(B)** Histogram of cell cycle time of daughter cells that divided during imaging (time to go from red cells to orange cells in pictorial representation of progeny number). **(C)** Histogram of distance moved by cells. **(D)** Histogram of radial displacement**. (E)** Histogram of angular displacement. Distance moved along the arc is considered as a proxy for angular displacement. **(F)** Angular displacement as a function of radial displacement. **(G)** Distance moved by cells as a function of their displacement.

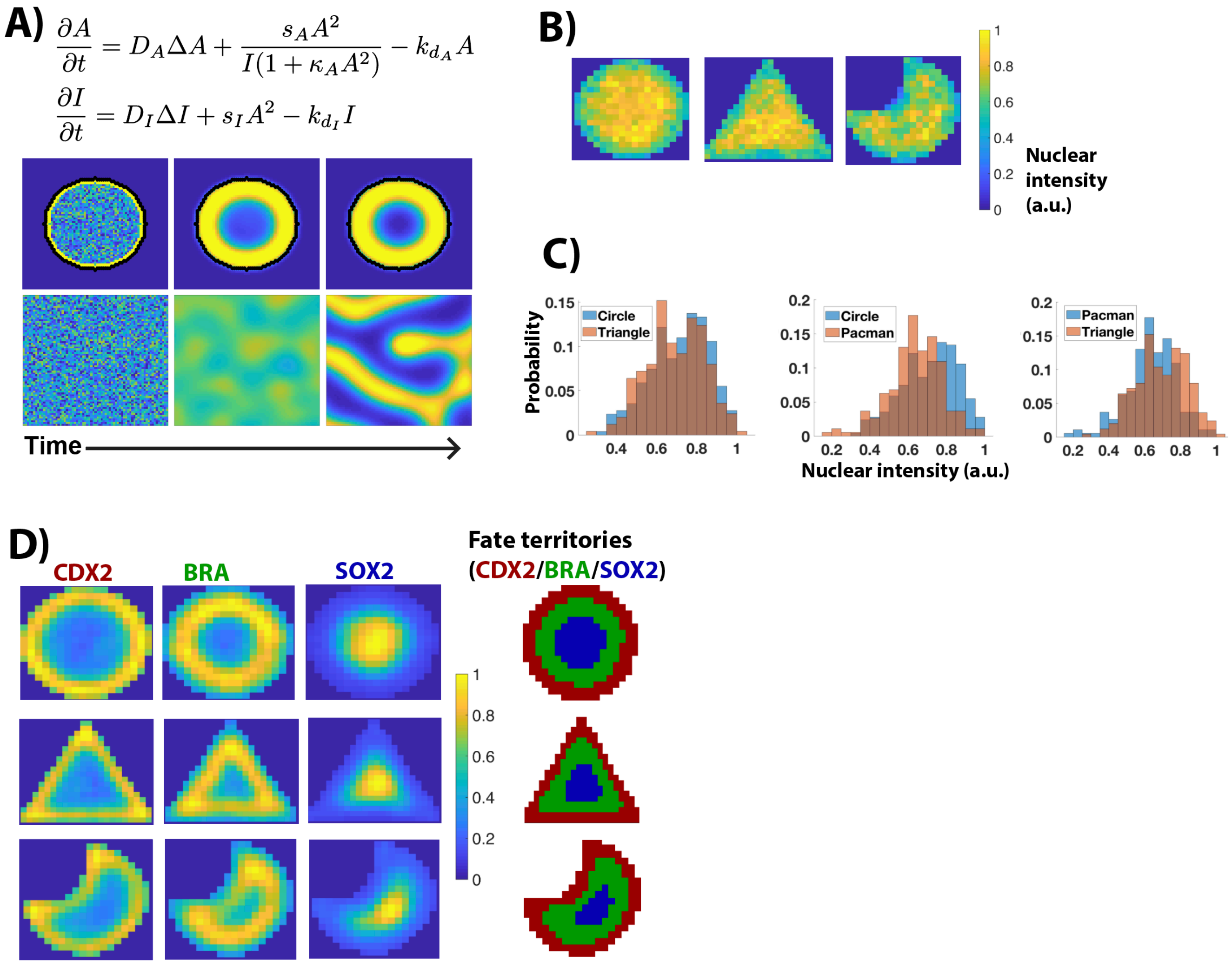


**Figure S6: Computing fate territories from experimental data, related to figure7**
**(A)** Equations and simulations for stripe-forming patterns. Simulation domain, assumptions and initial conditions are same as defined in Figure7. $D_A$ = 0.005, $D_I$ = 0.2, $s_A$ = 0.1, $s_I$ = 0.2, $kd_A$ = 0.1, $kd_I$ = 0.2, $\kappa_A$ = 0.25. degradation rate outside colony (kd = 0.5). **(B)** Nuclear maps, showing the nuclear intensity in each shape normalized to maximum nuclear intensity in circular colonies, at 44h post BMP treatment. **(C)** Histograms for A. As cell-seeding densities affect final fate patterns, we checked if average nuclear map for different shapes have comparable values. kstest2 returned 0 for nuclear maps of circle and triangle shapes, and 1 for nuclear maps of circle, pacman and triangle, pacman, indicating that nuclear map in pacman is different from the other two shapes. This is because the pacman has a higher perimeter/area ratio compared to the other two shapes and thus a higher fraction of regions with less nuclei (final cell density is lower at the colony edges). As the mean nuclear intensity for all three shapes was in the same order of magnitude (circle: 0.71, triangle: 0.69, pacman: 0.63), we considered them to be of comparable cell-seeding densities. **(D)** Intensity maps of indicated fate markers, normalized to the maximum intensity for the same marker in circular colonies. Fate territories were assigned by selecting the fate marker with the maximum intensity in that region. The nuclear and fate intensity maps represent values averaged over N=18 colonies in each of the three shapes. See also movie 14,15.

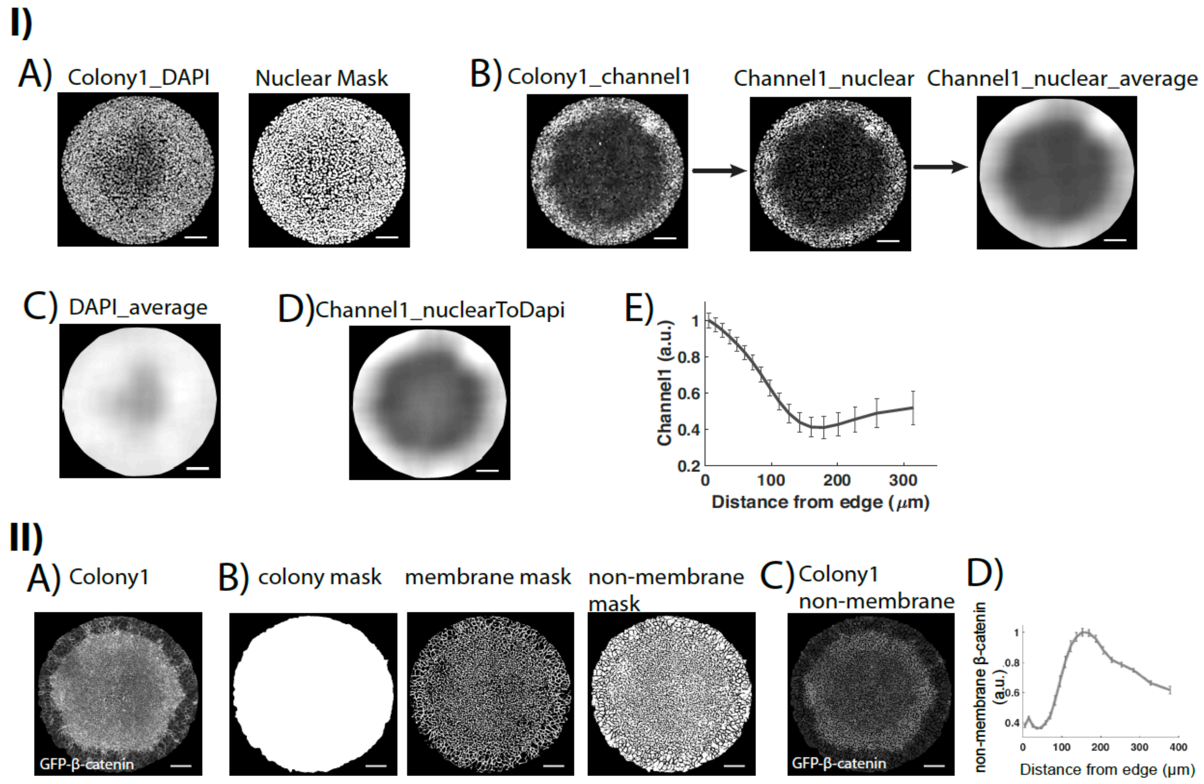


**S7: Analyses supplement**

All quantifications, plotted as intensity vs radial position (distance from colony edge) in the figures were created using one of the following two methods. ***(I)*** Channel1_nuclear. 1) For each colony, a nuclear mask was created by segmenting the DAPI image of that colony in Ilastik ((Sommer *et al.*, 2011) A). 2) The nuclear mask was then applied to each channel to extract nuclear pixels in that channel (B - Channel1_nuclear). 3) For each nuclear pixel in Channel1_nuclear, average local maxima were calculated in a region of radius 120um. (B – Channel1_nuclear_average). 4) Steps 2) and 3) were applied to DAPI image to get DAPI_Average (C). 5) Channel1_nuclear_average was normalized by DAPI_Average to get Channel1_nuclearToDAPI (D). 6) Mean intensity was calculated in different bins along the radius of Channel1_nuclearToDAPI to get radial averages. 7) Mean intensity in each bin was averaged across all colonies to get average radial average intensities (E). The final intensities were normalized to the intensity values in the same channel in the control sample (treated with BMP4 for 44h). Datasets quantified: Fate data (Figure1, 2D, 3F, 4G, 5A), SMAD2 data (Figure 3A, 4G). ***(II)*** GFP-β-Catenin_non-membrane. 1) For each colony, a membrane mask and a colony mask were created in Ilastik (Sommer *et al.*, 2011) using the GFP-β-catenin image (A, B). 2) Membrane mask was subtracted from colony mask to get non-membrane mask (B). 3) Non-membrane mask was applied to GFP-β-Catenin image to extract non-membrane β-catenin pixels (C). 4) As in *(I),* average intensity was calculated in different bins along the radius of Colony1_non-membrane images across different colonies. Datasets quantified: All GFP-β-catenin movies (Figure 2A, 3A, 4A). Error bars indicate standard error (D). Scale bar: 100μm

**Supplementary movies**

1) **Movie S1, Wnt signaling dynamics during fate patterning, related to Figure2**
GFP-b-catenin hESCs treated with 50ng/ml BMP4, imaged from 3-47h post treatment. Only non-membrane regions are shown. Colony diameter: 800um.

2) **Movie S2, Wnt signaling dynamics with LDN addition at 0h, related to Figure3B.**
GFP-b-catenin hESCs treated with 200nM LDN at 0h and 50ng/ml BMP4, imaged from 2-46h post treatment. Only non-membrane regions are shown. Colony size: 700um

3) **Movie S3, Wnt signaling dynamics with LDN addition at 11h, related to Figure3B.**
GFP-b-catenin hESCs treated with 50ng/ml BMP4, imaged from 2-46hpost treatment. 200nM LDN was added 11h post BMP treatment. Only non-membrane regions are shown. Colony size: 700um

4) **Movie S4, Wnt signaling dynamics with LDN addition at 23h, related to Figure3B.**
GFP-b-catenin hESCs treated with 50ng/ml BMP4, imaged from 2-46h post treatment. 200nM LDN was added 23h post BMP treatment. Only non-membrane regions are shown. Colony size: 700um.

5) **Movie S5, Wnt signaling dynamics with No LDN addition, related to Figure3B.**
GFP-b-catenin hESCs treated with 50ng/ml BMP4, imaged from 2-46h post treatment. No LDN was added in this sample. Only non-membrane regions are shown. Colony size: 700um.

6) **Movie S6, Wnt signaling dynamics with IWP2 addition at 15h, related to Figure4A.**
GFP-b-catenin hESCs treated with 50ng/ml BMP4 and 5μM IWP2, imaged from 2-47h post treatment. Only non-membrane regions are shown. Colony size: 700um.

7) **Movie S7, Wnt signaling dynamics with IWP2 addition at 30h, related to Figure4A.**
GFP-b-catenin hESCs treated with 50ng/ml BMP4, imaged from 2-47h post treatment. Only non-membrane regions are shown. Colony size: 700um

8) **Movie S8, Wnt signaling dynamics with no IWP2 addition, related to Figure4A.**
GFP-b-catenin hESCs treated with 50ng/ml BMP4, imaged from 2-46h post treatment. No IWP2 was added in this sample. Only non-membrane regions are

shown. Colony size: 700um.

9) **Movie S9, Cell movement during patterning, related to Figure6A.**
Cells with a nuclear fluorescent marker (RUES2-VENUS-H2B, green cells) mixed with unlabeled cells (ESI017) in the ratio 1:100, imaged from 20 to 47.5h post BMP treatment.

10) **Movie S10, Simulation of Activator-Inhibitor model in a circular colony, outside Turing regime, related to Figure7**

11) **Movie S11, Simulation of Activator-Inhibitor model in a lattice, outside Turing regime, related to Figure7**

12) **Movie S12, Simulation of Activator-Inhibitor model in a circular colony, within spot forming Turing regime, related to Figure7**

13) **Movie S13, Simulation of Activator-Inhibitor model in a lattice, within spot forming Turing regime, related to Figure7**

14) **Movie S14, Simulation of Activator-Inhibitor model in a circular colony, within stripe forming Turing regime, related to FigureS7**

15) **Movie S15, Simulation of Activator-Inhibitor model in a lattice, within stripe forming Turing regime, related to FigureS7**

### Mathematical modeling supplement

October 12, 2018

#### 1 Introduction

Our experimental results show that paracrine Wnt and Nodal signals move through spatially confined human embryonic stem cell colonies treated with BMP4 (gastruloids) to initiate self-organized fate patterning. The movement of these signals can be recapitulated by a simple reaction-diffusion based activator-inhibitor model defined below, where the activator(A)-inhibitor(I) pair corresponds to a composite of Wnt/Nodal and their secreted, diffusible feedback inhibitors.

$$\frac{\partial A}{\partial t} = D_A \Delta A + \frac{s_A A^2}{(k_I + I)} - k_{d_A} A \tag{1}$$

$$\frac{\partial I}{\partial t} = D_I \Delta I + s_I A^2 - k_{d_I} I \tag{2}$$

The equations are adapted from equations 1 in [2]. In sections2 and 3, we follow the stability analyses given in [2] to show that the parameter regime that recapitulates inward movement of signals takes the system to a homogeneous steady state and not a Turing pattern.

#### 2 Stability conditions

The qualitative behavior of the system of equations (1-2) depends on the choice of parameters. Although the possible parameter combinations are infinite, stability analyses can be used to determine parameter regimes corresponding to distinct system behaviors. Below, we derive conditions for the stability of the homogeneous steady state of the system and the formation spatial patterns.

To simplify analyses, we set $k_I + I = I'$. This transforms the equations to:

$$\frac{\partial A}{\partial t} = D_A \Delta A + \frac{s_A A^2}{I'} - k_{d_A} A \tag{3}$$

$$\frac{\partial I'}{\partial t} = D_I \Delta I' + s_I A^2 - k_{d_I} (I') + \sigma_I \tag{4}$$

where $\sigma_I = k_{d_I} * k_I$.

For simplicity, we assume that the system of equations operates in one dimensional closed domain [0,L], although the same analyses can be applied to a two-dimensional system. To non-dimensionalize the equations, we introduce new variables for length ($\bar{l}$), time ($\bar{t}$), and the concentrations ($\bar{A}$,$\bar{I}$) defined as:

$$\bar{t} = k_{d_A} * t$$
$$\bar{l} = (\sqrt{k_{d_A}/D_I}) * l$$
$$\bar{A} = \frac{k_{d_A} * s_I}{k_{D_I} * s_A} * A$$
$$\bar{I} = \frac{k_{d_A}^2 * s_I}{k_{D_I} * s_A^2} * I'$$

Rewriting equations (3),(4) with new variables yields:

$$\frac{\partial \bar{A}}{\partial \bar{t}} = D\bar{\Delta}\bar{A} + \frac{\bar{A}^2}{\bar{I}} - \bar{A} \tag{5}$$
$$\frac{\partial \bar{I}'}{\partial \bar{t}} = \bar{\Delta}\bar{I}' + \mu(\bar{A}^2 - \bar{I}) + \sigma \tag{6}$$

where $\sigma = \frac{k_I * k_{D_I} * s_I}{s_A^2}$ , $\mu = \frac{k_{D_I}}{k_{D_A}}$, $D = \frac{D_A}{D_I}$, $\bar{\Delta} = \partial^2/\partial\bar{x}^2$

In the absence of diffusion, the system has three steady states, which are given by:
$A_0 = 0$, $I_0 = \frac{\sigma}{\mu}$
$A_0 = I_0 = \frac{\mu \pm \sqrt{\mu^2 - 4\mu\sigma}}{2\mu}$
These correspond to two stable points (at $A_0 = 0$ and $A_0 = \frac{\mu + \sqrt{\mu^2 - 4\mu\sigma}}{2\mu}$, with a saddle point at $A_0 = \frac{\mu - \sqrt{\mu^2 - 4\mu\sigma}}{2\mu}$. All the subsequent analyses are performed for the stable steady state with non-zero $A_0$.

To determine the stability of the steady state, we introduce a small perturbation ($|\delta A_0|, |\delta I_0| << 1$) around steady state:

$$\bar{A} = A_0 + \delta A \tag{7}$$
$$\bar{I}' = I_0 + \delta I \tag{8}$$

where

$$\delta A = \delta A_0 e^{\omega t} cos(kx) \tag{9}$$
$$\delta I = \delta I_0 e^{\omega t} cos(kx) \tag{10}$$

k is the wavenumber associated with the spatial perturbation. The system operates in a closed domain ([0,L]) with boundary conditions such that the flux vanishes at the boundaries:

$$\left.\frac{\partial A}{\partial x}\right|_{x=0,x=L} = \left.\frac{\partial I}{\partial x}\right|_{x=0,x=L} = 0 \tag{11}$$

Thus, k takes only discrete values: $k_n = n\pi/L$, n = 0,1,2...

Introducing values from (7-10) in (5-6), linearizing the reaction terms and retaining terms up to first order, gives:

$$\begin{pmatrix} \omega + k^2 D - 1 & 1 \\ -2\mu A_0 & \omega + k^2 + \mu \end{pmatrix} \begin{pmatrix} \delta A_0 \\ \delta I_0 \end{pmatrix} = 0 \tag{12}$$

The perturbation amplitudes($\delta A_0, \delta I_0$) are different from zero if and only if,

$$\begin{vmatrix} \omega + k^2 D - 1 & 1 \\ -2\mu A_0 & \omega + k^2 + \mu \end{vmatrix} = 0 \tag{13}$$

which implies,

$$\omega^2 + \alpha\omega + \beta = 0 \tag{14}$$

$$\omega = \frac{-\alpha \pm \sqrt{(\alpha^2 - 4\beta)}}{2}$$

where

$$\alpha = k^2(D+1) + \mu - 1 \tag{15}$$
$$\beta = Dk^4 + (\mu D - 1)k^2 + \mu(2A_0 - 1) \tag{16}$$

To obtain a diffusion driven instability, it is necessary that the perturbation grows with time i.e. $\mathrm{Re}[\omega(k)] > 0$, and the wavelength of the pattern fits in the spatial domain. Assuming that the the length 'L' of the domain is much larger than the wavelength of the pattern, we can derive conditions for spatial patterning. In this length regime, we consider the wavenumber 'k' as a continuous variable greater than 0. Perturbations grow at this wavenumber if $\mathrm{Re}[\omega(k)] > 0$. This happens if one of the following two conditions is true:
1) $\alpha < 0$
The condition $\alpha < 0$ equates to

$$k^2(D+1) + \mu < 1 \tag{17}$$

As k, D and $\mu$ are positive constants, $\mu < k^2(D+1) + \mu$ . Thus,

$$\mu < k^2(D+1) + \mu < 1 \tag{18}$$

If $\mu < 1$, there exist values of k such that inequality (17) is satisfied. This instability can be achieved without invoking diffusion, and thus, is not diffusion driven and does not result in any spatial pattern.

2) $\alpha > 0$ and $\beta < 0$
By the same reasoning as above, requiring $\alpha > 0$ for all k $\in$ R yields,

$$\mu > 1 \tag{19}$$

The condition $\beta < 0$ equates to

$$Dk^4 + (\mu D - 1)k^2 + \mu(2A_0 - 1) < 0 \tag{20}$$

The left hand side of the above inequality represents a concave parabola in the variable $\gamma = k^2$.

$$f(\gamma) = D\gamma^2 + (\mu D - 1)\gamma + \mu(2A_0 - 1) \tag{21}$$

$f(\gamma) < 0 \implies \min(f(\gamma)) < 0$. This minimum of $f(\gamma)$ is given by the value of $\gamma$ for which

$$\begin{gathered} \frac{df}{d\gamma} = 0 \\ 2D\gamma + \mu D - 1 = 0 \\ \gamma = \frac{1 - \mu D}{2D} \end{gathered} \tag{22}$$

As $\gamma = k^2$ and $k^2 > 0$, from equation (22), we get

$$\mu D < 1 \tag{23}$$

Now, $\min(f(\gamma)) < 0$ gives

$$D(\frac{1-\mu D}{2D})^2 + (\mu D - 1)(\frac{1-\mu D}{2D}) + \mu(2A_0 - 1) < 0 \tag{24}$$

Simplifying equation (24) gives

$$\mu < \frac{(\sqrt{2A_0} - \sqrt{2A_0 - 1})^2}{D} \tag{25}$$

From (19) and (25),

$$1 < \mu < \frac{(\sqrt{2A_0} - \sqrt{2A_0 - 1})^2}{D} \tag{26}$$

This instability occurs only in the presence of diffusion, and is therefore diffusion driven. Within some finite range of wavenumbers k (k > 0), this instability will form spatial pattern provided the length of reaction domain is large enough to fit the spatial perturbations.

The homogeneous steady state would be stable if $\alpha > 0$ and $\beta > 0$, which, from above analyses equates to:

$$\mu > \frac{(\sqrt{2A_0} - \sqrt{2A_0 - 1})^2}{D} \tag{27}$$

To summarize,
1) A diffusion driven instability occurs if and only if $\mu > 1$ & $D < \frac{(\sqrt{2A_0} - \sqrt{2A_0 - 1})^2}{\mu}$
2) System reaches a homogeneous steady state if $\mu > 1$ & $D > \frac{(\sqrt{2A_0} - \sqrt{2A_0 - 1})^2}{\mu}$

(It must be noted that $A_0$ is a function of $\mu$ and $\sigma$.)

### 3 Simulation components

#### 3.1 Simulation domain

All simulations were performed on a 2d lattice of size 190 by 190 pixels. To incorporate a circular colony, a circle of radius 25 pixels was defined at the center of the lattice. Experimentally, the circle is analogous to a circular colony of radius 400$\mu$m, and the lattice region outside the circle is analogous to media. $\therefore$ 1 pixel = 16$\mu$m.

#### 3.2 Model parameters

The time step for running the simulation(dt) was 0.1. Assuming the unit of time is minutes, dt = 0.1min and the unit of concentration is picomoles(pM). With 1 pixel = 16$\mu$m and 1 time step = 0.1min, the parameter values that recapitulate inwards movement of paracrine signaling activities are as follows:

These parameter values are in the same order of magnitude as observed experimentally [3], [1].

#### 3.3 Turing patterns

Keeping all the parameters unchanged, transition into Turing pattern formation regime is possible by reducing the diffusion constant of the activator ($D_A$) (equation 26). For the above mentioned parameter set, the critical value of $D_A$, at which the system bifurcates is $0.0087 pixel^2/(0.1 min) = 0.3712 \mu m^2/s$. It must be noted that near the bifurcation point, even within the Turing regime, the activator concentration in the lattice appears to be homogeneous as the difference between values across the lattice is negligible.

| Parameters | Values in simulation | Values with units |
|---|---|---|
| $D_A$ | 0.02 | $0.85 \mu m^2/s$ |
| $D_I$ | 0.4 | $17 \mu m^2/s$ |
| $k_{d_A}$ | 0.001 | $1.6 * 10^{-4}/s$ |
| $k_{d_I}$ | 0.008 | $12.8 * 10^{-4}/s$ |
| $s_A$ | 0.01 | $1.6 * 10^{-3}/s$ |
| $s_I$ | 0.01 | $1.6 * 10^{-3}/(s)$(pM) |
| $k_I$ | 1 | 1pM |

### References


[1] Uri Alon. *An introduction to systems biology: design principles of biological circuits.* Chapman and Hall/CRC, 2006.

[2] AJ Koch and Hans Meinhardt. Biological pattern formation: from basic mechanisms to complex structures. *Reviews of modern physics*, 66(4):1481, 1994.

[3] Patrick Müller, Katherine W Rogers, Ben M Jordan, Joon S Lee, Drew Robson, Sharad Ramanathan, and Alexander F Schier. Differential diffusivity of nodal and lefty underlies a reaction-diffusion patterning system. *Science*, 336(6082):721–724, 2012.
